# LDL1 and LDL2 histone demethylases interact with FVE to regulate flowering in *Arabidopsis*

**DOI:** 10.1101/2022.05.12.491658

**Authors:** Mahima, Sourav Chatterjee, Sharmila Singh, Ananda K. Sarkar

## Abstract

In higher plants, epigenetic modifications provide a stage for both transient and permanent cellular reprogramming required for vegetative to reproductive phase transition. *Arabidopsis* LSD1-like 1 (LDL1), a histone demethylase positively regulates floral transition, but the molecular and biochemical nature of LDL1 mediated flowering is poorly understood. Here we have shown that LDL1 mediated regulation of flowering is dependent on MADS AFFECTING FLOWERING 4 (MAF4) and MAF5 floral repressors. LDL1 binds on the chromatin of *MAF4* and *MAF5* and removes H3K4me2 activation marks to repress their expression. Further we show that LDL2 negatively regulates the expression of *MAF4* and *MAF5* redundantly with LDL1. Both LDL1 and LDL2 interact with an autonomous flowering pathway protein, FLOWERING LOCUS VE (FVE), to regulate the floral transition and thus could be a part of the FVE-corepressor complex. We show that MAF5 interacts with other floral repressors FLC and SHORT VEGETATIVE PHASE (SVP) and repress the expression of *FT* to delay floral transition. Thus, our results deepen the mechanistic understanding of LDL1/LDL2-FVE mediated floral transition in *Arabidopsis*.

## Introduction

In plants, the precise timing of the transition from the vegetative to reproductive phase is crucial for deciding reproductive success ^1^. *Arabidopsis* has six major genetic pathways which coalesce various internal and external signals to access the appropriate time of flowering. These pathways include autonomous, photoperiod, vernalization, gibberellin (GA), temperature, and age-dependent pathways ^2–4^. All these pathways either converge to suppress the expression of MADS-box transcription factor, *FLOWERING LOCUS C* (*FLC*), or directly upregulate the floral integrator genes, *FLOWERING LOCUS T* (*FT*) and *SUPPRESSOR OF OVEREXPRESSION OF CONSTANS1* (*SOC1*), which are negatively regulated by FLC ^5–7^. FLC associates with another MADS-box protein, SHORT VEGETATIVE PHASE (SVP), and together they repress *FT* in companion cells of leaf vein and *SOC1* in shoot meristem ^7–9^. Lately, the role of FLC clade members, MADS AFFECTING FLOWERING1 to 5 (MAF1-to 5) has been implicated in preventing precocious flowering. MAF1 was the first FLC homolog to be analyzed, and it seems to act through both photoperiod and thermosensory pathways in parts, independently of FLC ^10–12^. *MAF2* to 4 are also reported to delay flowering redundantly with FLC ^13–15^. MADS-box proteins are known to form multimeric protein complexes in different combinations ^16, 17^. FLC, SVP, FLM, and MAF3 are reported to be highly enriched in intronic regions of *FT* chromatin, and MAF2 and MAF4 are predicted to bind *FT* chromatin as well ^8, 9, 15^. Therefore, it is possible that SVP forms dimeric/tetrameric MADS domain repressor complexes with different combinations of FLC clade proteins to regulate flowering. All these MADS-box proteins bind to CArG motifs enriched in the intronic and 5’ promoter region of the *FT* locus, possibly with different affinities and/or tissue preferences to repress *FT* expression in a partially overlapping manner ^7, 15, 18, 19^.

Epigenetic modifications play a crucial role in negatively regulating *FLC* and its clade members during vernalization and in the autonomous flowering pathway. In winter annual ecotypes of *Arabidopsis*, vernalization, a prolonged cold exposure, is a prerequisite to initiating flowering. Vernalization replaces activating H3K4me3 and H3K36me3 modifications with repressive H3K27me3 modification via the activity of various chromatin modifiers ^20^. Similarly, in the autonomous flowering pathway, various epigenetic modifiers act together to repress the expression of *FLC* ^21^. One of the corepressor complexes of the autonomous flowering pathway includes FLOWERING LOCUS D (FLD), FLOWERING LOCUS VE (FVE), HISTONE DEACETYLASE 5 (HDA5), and HDA6. This complex removes activating H3K4 methylation and H3 or H4 acetylation marks from *FLC, MAF4*, and *MAF5* loci ^22–26^. FLD belongs to the LSD1-like (LDL) family of histone demethylases in *Arabidopsis*, which includes LDL1, LDL2, and LDL3. These proteins are homologs of human LYSINE SPECIFIC DEMETHYLASE 1 (LSD1) ^27^. LSD1 contains an N-terminal Swi3p/Rsc8p/Moira (SWIRM) domain involved in protein-protein interaction and a C-terminal amine-oxidase like (AOL) domain ^28, 29^. The AOL domain further contains two subdomains, a FAD-binding, and a substrate-binding domain ^30, 31^. Like LSD1, *Arabidopsis* LDL family members also comprise an N-terminal SWIRM domain and a C-terminal amine oxidase domain ^27^. LDL1 and LDL2 suppress various seed dormancy-related genes, like *DELAY OF GERMINATION 1* (*DOG1*), and genes related to Abscisic acid biosynthesis and signaling ^32^. A recent report has elucidated the role of LDL1 and LDL2 in regulating circadian rhythm, where they negatively regulate the expression of the evening expressed *TIMING OF CAB EXPRESSION 1* (*TOC1*) ^33^. Additionally, LDL1 also inhibits root growth and branching by negatively regulating the expression of *LATERAL ROOT PRIMORDIUM1* (*LRP1*) in the primary root and is proposed to suppress root branching by modulating the expression of *AUXIN RESPONSE FACTORS* (*ARFs*) ^34, 35^.

The two mutant alleles of *LDL1, ldl1-1* and *ldl1-2* show late flowering phenotype ^27, 36^. This late flowering phenotype indicates the potential role of LDL1 in the regulation of flowering time, which is underexplored. In the present study we have shown the mechanism behind LDL1 mediated flowering time regulation by identifying its novel downstream targets and biochemical activity. We have shown that LDL1 and LDL2 targets the same set of MADS-box floral repressors to allow flowering and interact with FVE, a crucial component of the autonomous flowering pathway. MAF5 a common target of LDL1 and LDL2 was found to interact with other floral repressors, FLC and SVP, and subsequently repress the expression of *FT*. Collectively, these findings enhance our understanding on the role of chromatin modifiers in the regulation of flowering time.

## Results

### LDL1 negatively regulates the expression of *MAF4 and MAF5*

LDL1 belongs to the LDL family of histone demethylases in *Arabidopsis*. LDL1, along with its family members, LDL2 and FLD allow vegetative to floral transition by negatively regulating the expression of *FLC* ^27, 36^. To understand the genetic interaction between *LDL1* and *FLC*, we generated a *ldl1flc* double mutant. We observed that *ldl1flc* plants flowered earlier than *ldl1* but later than *flc* plants, suggesting that there might be other targets of LDL1 other than *FLC* (Supplementary Figure 1). FLD, a family member of LDL1 regulates the expression of two other members of MADS-box family genes, *MAF4* and *MAF5* ^24^. This prompted us to check the expression of *MAF4* and *MAF5* in *ldl1* mutant plants and we found that the expression of *FLC, MAF4*, and *MAF5* transcripts was upregulated in *ldl1* with respect to the wild type (WT) (Figure 1A)

**Figure 1.**
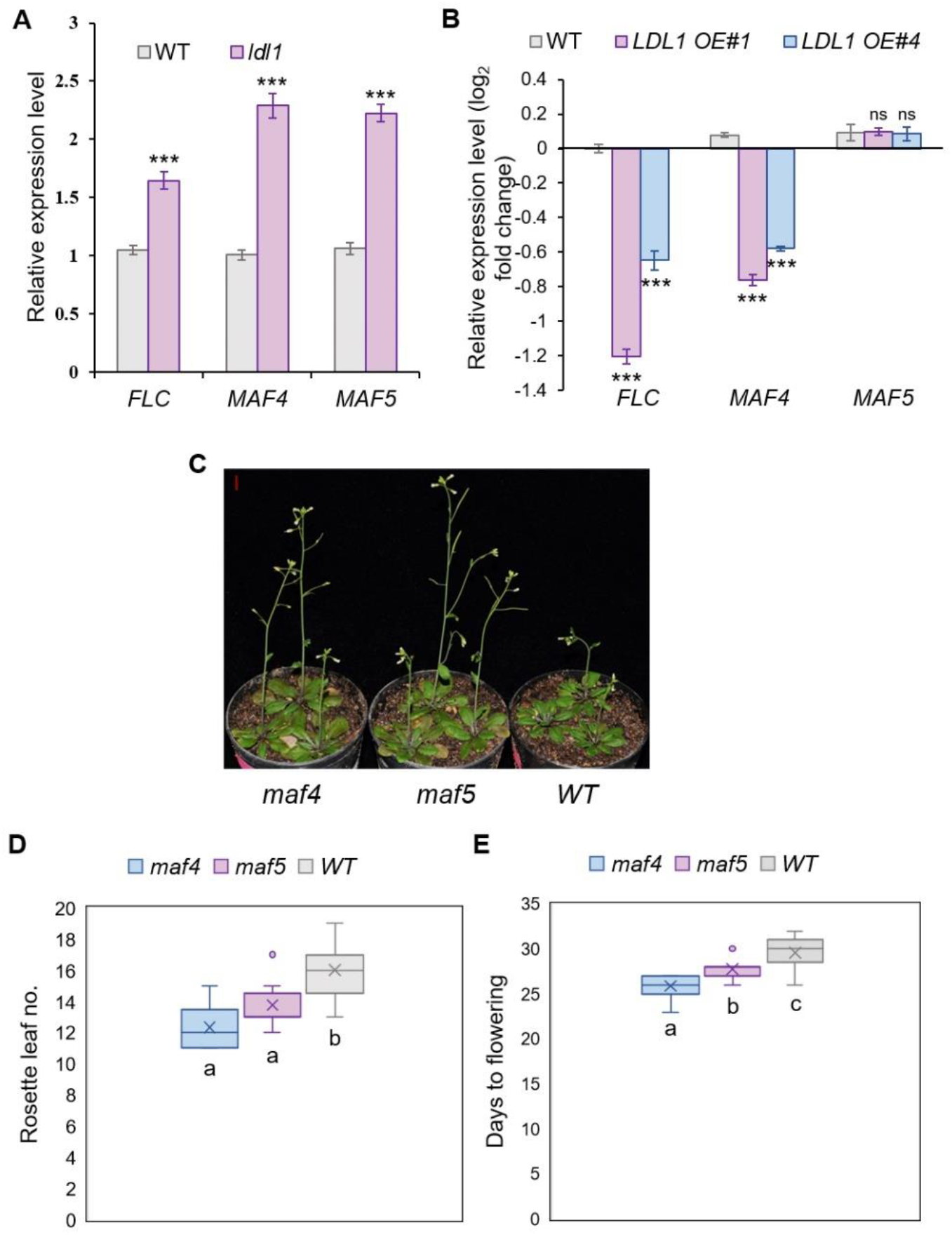
LDL1 promotes flowering by negatively regulating the expression of *MAF4* and *MAF5*. (**A**) Relative expression of *FLC, MAF4*, and *MAF5* in WT and *ldl1*. (**B**) Relative expression of *FLC, MAF4*, and *MAF5* in WT, *LDL1 OE#1*, and *LDL1 OE#4*. (**C**) Flowering phenotype of *maf4*, and *maf5* with respect to WT plants under long-day conditions. Both *maf4* and *maf5* mutant plants show earlier flowering than WT plants. (**D**) Rosette leaf numbers of *maf4, maf5*, and WT plants at bolting (n=15). (**E**) Days taken to flower by *maf4, maf5*, and WT plants (n=15). Expression of *MAF4* and *MAF5* was checked in the shoots of 14 days old seedlings. Error bars indicate the standard error (± SE) of three independent experiments. Asterisks indicate significant differences (\**p* ≤ 0.05, \*\**p* ≤ 0.01, \*\*\**p* ≤ 0.001; unpaired two tailed *t-*test) in (**A**) & (**B**). Scale bar=1cm in (**C**). Different letters on whiskers of box plots in (**D**) & (**E**) indicate statistically significant differences (one-way ANOVA followed by post-hoc Tukey’s test, p <0.05).

To understand the role of LDL1 more elaborately in relation to the regulation of *MAF4* and *MAF5* expression, we generated the overexpression construct of *LDL1* under constitutive RPS5A promoter (*proRPS5A:LDL1*) ^37^. Several T1 plants showed upregulation of *LDL1* transcripts, and we selected two independent T1 plants which showed the highest upregulation, *LDL1 OE#1* and *LDL1 OE#4* (Supplementary Figure 2). We selected homozygous T3 plants and checked the expression of *FLC, MAF4*, and *MAF5* transcripts in *LDL1 OE#1* and *LDL1 OE#4* plants. Both *FLC* and *MAF4* were downregulated in the *LDL1 OE* plants, whereas transcript levels of *MAF5* were comparable to the WT plants (Figure 1B).

To confirm that LDL1 regulated floral transition is dependent on MAF4 and MAF5, we proceeded to check whether *maf4* and *maf5* single mutants have altered flowering time. As previously reported, *maf4* and *maf5* show early floral transition in the *Landsberg erecta* (*ler*) ecotype, but their role as floral repressors in *Columbia* (Col-0) ecotype remained uncertain ^13^. We measured the days to bolting in *maf4* and *maf5* single mutants and found that both *maf4* and *maf5* plants showed early flowering phenotype as compared to the WT plants (Figure 1C). Rosette leaf numbers were also in accordance with the time taken to flower (Figure 1D and 1E). This result indicates that both MAF4 and MAF5 are involved in negatively affecting flowering time. Taken together, our results specify that LDL1 induces floral transition through repressing *FLC, MAF4*, and *MAF5*.

### LDL1 binds to the chromatin of *MAF4* and *MAF5* to regulate their expression

LSD1, the human homolog of LDL1 regulates its targets by binding to their chromatin and carrying out histone demethylation ^38^. To check whether LDL1 binds to the chromatin of *MAF4* and *MAF5* directly to regulate their expression, we generated a translational fusion construct of LDL1 with β-glucuronidase (GUS) under its native promoter (*proLDL1:LDL1-GUS)*. To confirm that the *proLDL1:LDL1-GUS* construct is functional, we transformed the construct into *ldl1* mutant background. The level of *LDL1* mRNA was restored and the late-flowering phenotype of *ldl1* mutant plants was rescued in *proLDL1:LDL1-GUS (ldl1)* plants (Supplementary Figure 3). Through Histochemical GUS assay, LDL1 was found to be expressed in shoot apical meristem, leaves, flowers, hypocotyl, primary root, and different stages of lateral root (LR) development, indicating its potential role in various aspects of plant development (Supplementary Figure 4).

Using *proLDL1:LDL1-GUS* (*ldl1*) plants, we checked the binding of LDL1 on the chromatin of *MAF4* and *MAF5* through ChIP-quantitative Real-Time (ChiP-qRT) PCR. We found the enrichment of LDL1 on the promoter and exon1 of *MAF4* and *MAF5* (Figure 2A and 2B). Next, we generated the reporter constructs of *MAF4* and *MAF5* to see how their expression patterns differ in *ldl1* mutant plants from the WT plants. We transformed the *proMAF4:GFP-GUS* and *proMAF5:GFP-GUS* constructs into the *ldl1* mutant and WT plants (Figure 2C). We selected the T1 plants which showed the GUS staining in both *ldl1* and WT backgrounds for performing further experiments. In the T2 generation, the GUS activity of *proMAF4:GFP-GUS* was observed in the hypocotyl and leaves of 7-dag seedlings in the *ldl1* background but was absent in the WT background (Figure 2D and 2E). Similarly, for *proMAF5:GFP-GUS*, we found the GUS activity in the shoot of 9dag seedlings of *ldl1* background, which was absent in the WT background (Figure 2F and 2G). T2 lines obtained from four independent T1 lines of each construct in both WT and *ldl1* backgrounds were checked for GUS staining, and the results were consistent. Collectively, our data suggest that LDL1 binds to the chromatin of *MAF4* and *MAF5* and negatively regulates their expression *in planta*.

**Figure 2.**
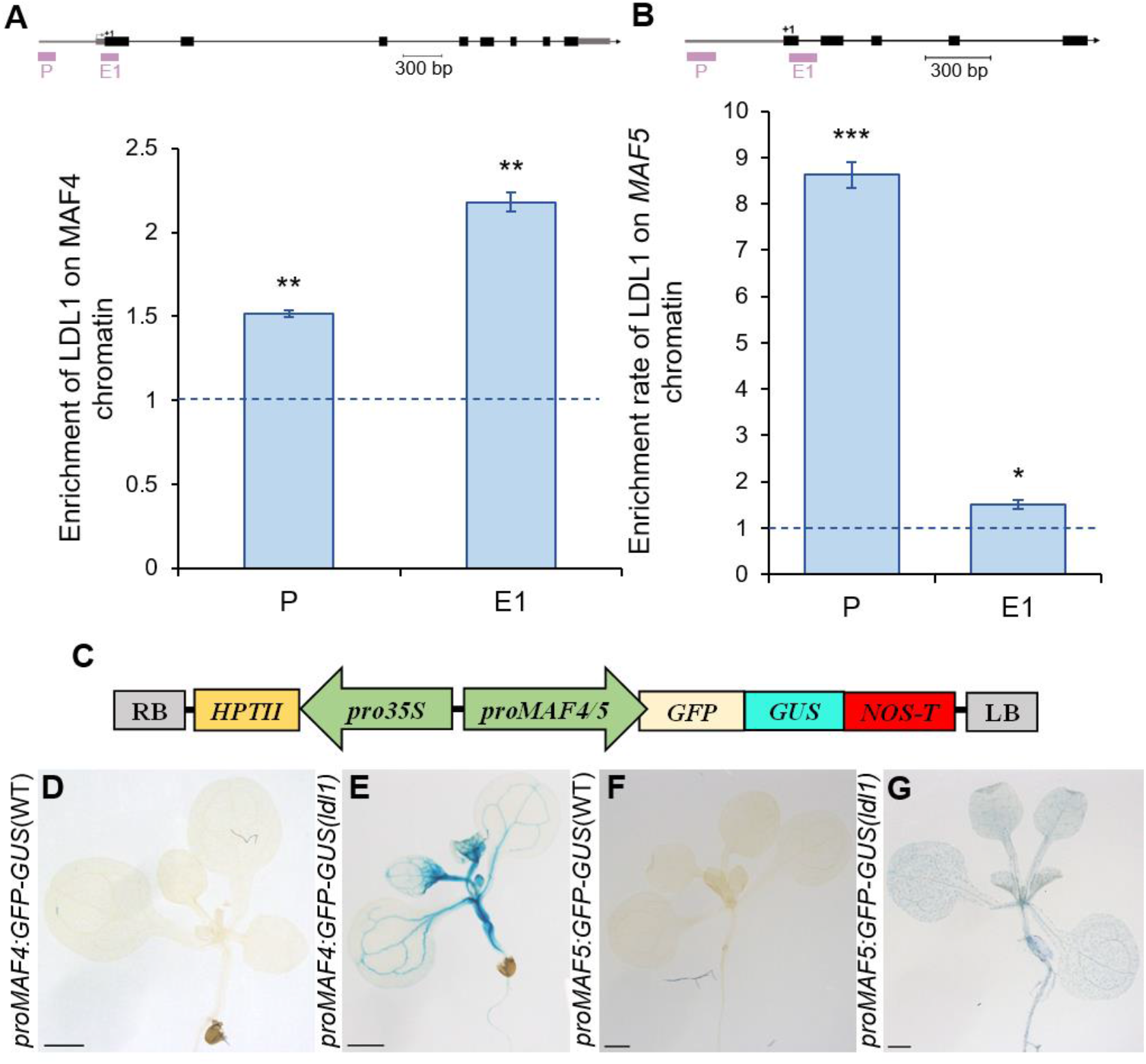
LDL1 directly binds to the chromatin of *MAF4* and *MAF5* to regulate their expression. (**A**) Enrichment of LDL1 on the promoter and 1st exon of *MAF4* chromatin. (**B**) Enrichment of LDL1 on the promoter and 1st exon of *MAF5* chromatin. (**C**) Construct map of *proMAF4/proMAF5:GFP-GUS*. (**D & E**) Expression of *proMAF4:GFP-GUS* in WT and *ldl1* background at 7dag. (**F & G**) Expression of *proMAF5:GFP-GUS* in WT and *ldl1* background at 9dag. Error bars indicate the standard error (± SE) of three independent experiments. Asterisks indicate significant differences (\**p* ≤ 0.05, \*\**p* ≤ 0.01, \*\*\**p* ≤ 0.001; unpaired two tailed *t-*test). Scale bar=1mm in (**D**), (**E**), (**F**) & (**G**).

### Mutations in *MAF4* and *MAF5* loci rescue the late-flowering phenotype of *ldl1*

To understand the interaction between LDL1 and MAF4 and MAF5 at a genetic level, we crossed *ldl1* with *maf4* and *maf5* single mutants and selected *ldl1maf4* and *ldl1maf5* double mutant plants. We found that both *ldl1maf4* and *ldl1maf5* plants flowered earlier than the *ldl1* single mutant plan and their flowering time was comparable to that of the WT plants (Figure 3A and 3D). Rosette leaves quantification of *ldl1, ldl1maf4, ldl1maf5*, and WT plants at the time of bolting was consistent with their flowering phenotype, that is, the number of rosettes leaves of *ldl1maf4* and *ldl1maf5* during bolting was comparable to that of the WT plants, whereas *ldl1* single mutant plants had more rosette leaves in accordance with its late-flowering phenotype (Figure 3B, 3C, 3E, and 3F). Therefore, our results indicate LDL1 mediated flowering time regulation is dependent on MAF4 and MAF5 functions.

**Figure 3.**
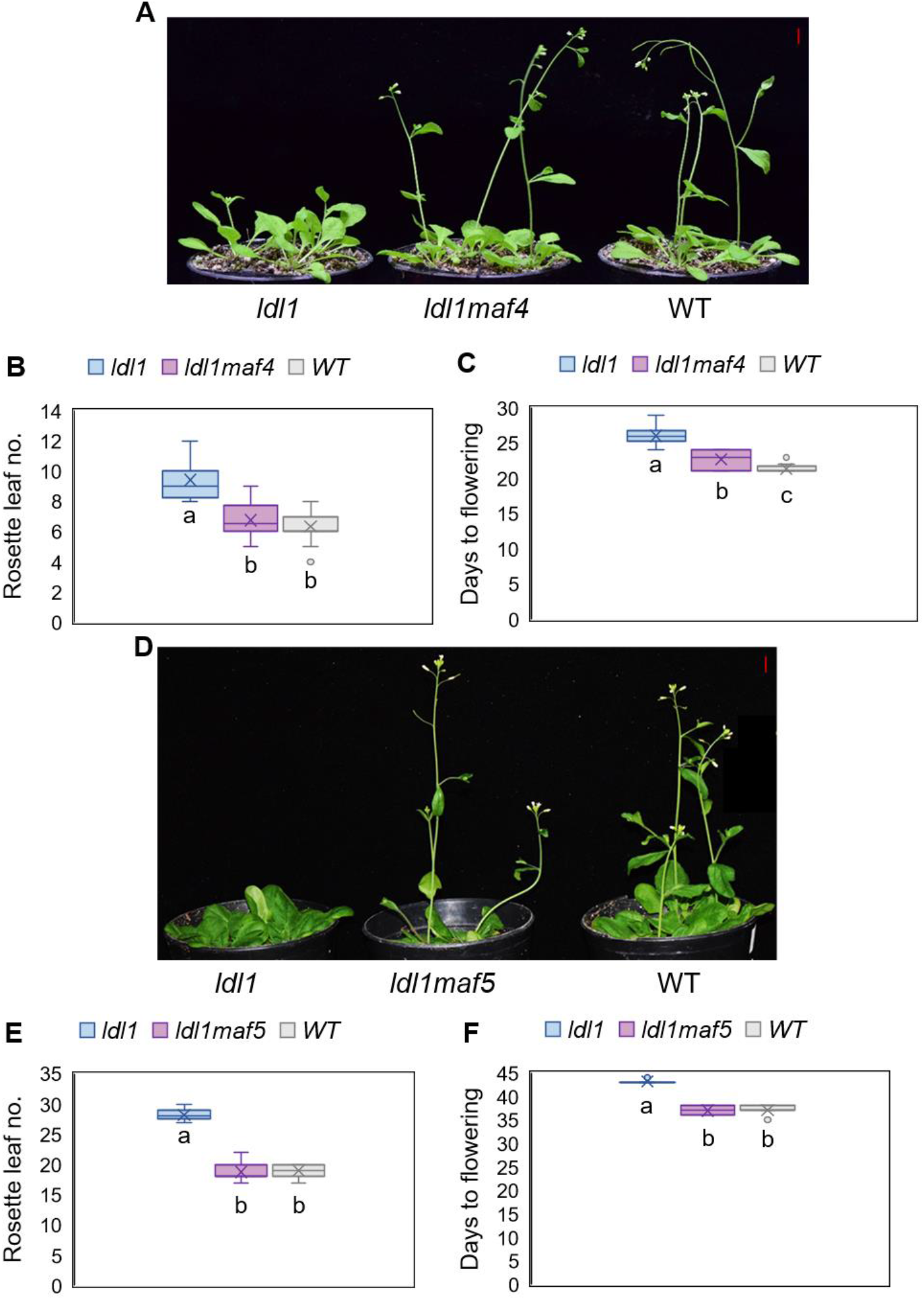
Mutation in *MAF4* and *MAF5* rescues the late flowering phenotype of *ldl1*. (**A**) Flowering phenotype of *ldl1maf4* with respect to *ldl1* and WT plants. (**B**) Rosette leaf numbers of *ldl1, ldl1maf4* and WT plants at bolting (n=15). (**C**) Days taken to flower by *ldl1, ldl1maf4*, and WT plants (n=15). (**D**) Flowering phenotype of *ldl1maf5* with respect to *ldl1* and WT. (**E**) Rosette leaf numbers of *ldl1, ldl1maf5*, and WT plants at bolting (n=15). (**F**) Days taken to flower by *ldl1, ldl1maf5*, and WT (n=15). Different letters on whiskers of box plots indicate statistically significant differences (one-way ANOVA followed by post-hoc Tukey’s test, p <0.05). Scale bar=1cm in (**A**) and (**D**).

### LDL1 shows H3K4me2 and H3K9me2 demethylase activity in vitro

The human homolog of LDL1, LSD1 majorly acts as a transcriptional repressor by removing activating histone marks, H3K4me1 and H3K4me2 by flavin adenosine dinucleotide (FAD) dependent oxidation reaction, but an interaction of LSD1 with androgen receptor results in H3K9 demethylation leading to gene activation ^39, 40^. In comparison to its human counterpart, biochemical nature of LDL1 is poorly understood ^27, 36^. Therefore, to check the biochemical activity of LDL1 and understand how it represses *MAF4* and *MAF5*, we purified GST tagged LDL1 and checked its demethylation activity on histones (Figure 4A). We found that like LSD1, *Arabidopsis* LDL1 too has *in-vitro* H3K4me2 demethylase activity (Figure 4B and 4C). In addition, LDL1 was able to demethylate H3K9me2 marks *in-vitro* (Figure 4B and 4D), which is not the case for LSD1 as it is unable to demethylate H3K9 marks *invitro* and requires the specific interacting partners to carry out H3K9 demethylation ^40, 41^. Taken together with previous results, we can deduce that LDL1 binds to the chromatin of *MAF4* and *MAF5* and repress them by removing activating H3K4me2 marks.

**Figure 4.**
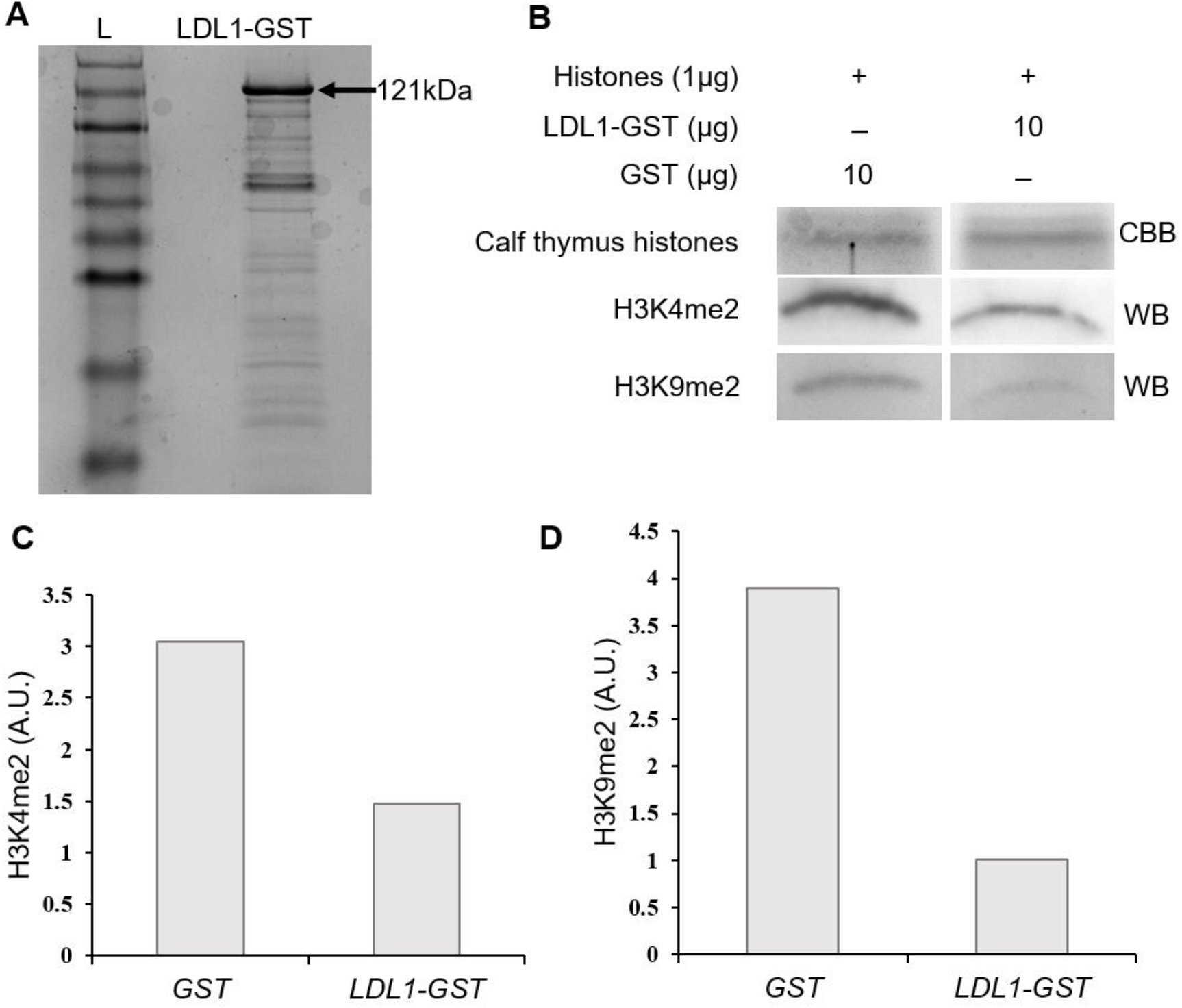
LDL1 has invitro H3K4me2 and H3K9me2 demethylase activity. (**A**) LDL1-GST after purification and concentration. (**B**) LDL1 demethylation assay followed by western blot using H3K4me2 and H3K9me2 specific antibodies. (**C**) and (**D**) Quantification of bands obtained by western blotting by ‘imageJ’ software. A.U.=arbitrary units.

### LDL2 along with LDL1 negatively regulates the expression of MAF4 and MAF5

Since LDL family members FLD and LDL1 regulate flowering, we were interested to know if LDL2 also affects flowering. To understand this, we scored the time taken for bolting by *ldl2* plants. Like *ldl1* plants, *ldl2* plants have a late-flowering phenotype compared to WT, but it is not as strong as that of *ldl1* plants (Figure 5A and 5B). The *ldl1ldl2* plants have a stronger late-flowering phenotype than either of the single mutants (Figure 5A and 5B). LDL2 is also known to negatively regulates the expression of *FLC* and *FWA* along with LDL1 ^27^. Given the flowering phenotype of *ldl2* and previous data, we were interested to find whether LDL2 regulates the expression of *MAF4* and *MAF5*. The expression of both *MAF4* and *MAF5* was upregulated in *ldl1*and *ldl2* plants in comparison to WT plants. The expression of *MAF4* and *MAF5* in *ldl1ldl2* plants was even higher than either of the single mutant plants which agrees with their flowering phenotype (Figure 5A to 5D). Taken together, our phenotypic and expression analysis suggest that LDL1 and LDL2 regulate flowering synergistically.

**Figure 5.**
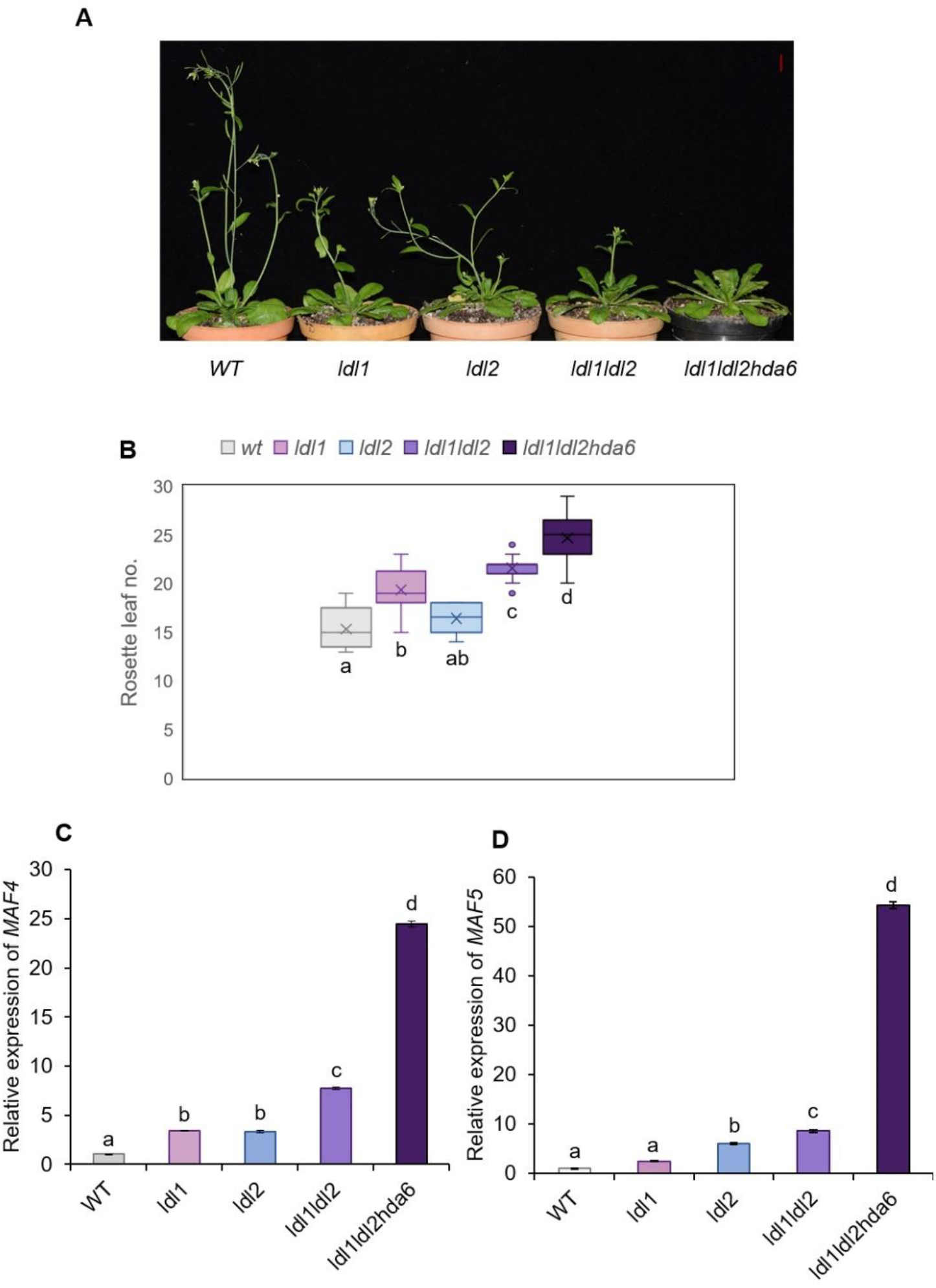
LDL1, along with LDL2 and HDA6, regulates the expression of *MAF4* and *MAF5*. Flowering phenotype of *ldl1ldl2* and *ldl1ldl2hda6* with respect to *ldl1, ldl2*, and WT plants. Rosette leaf numbers of WT, *ldl1*, l*dl2, ldl1ldl2*, and *ldl1ldl2hda6* plants at bolting (n=15). (**C**) Relative expression of *MAF4* in WT, *ldl1, ldl2, ldl1ldl2*, and *ldl1ldl2hda6*. (**D**) Relative expression of *MAF5* in WT, *ldl1, ldl2, ldl1ldl2*, and *ldl1ldl2hda6*. Expression of *MAF4* and *MAF5* was checked in the shoots of 14 days old seedlings. Error bars indicate the standard error (± SE) of three independent experiments. different letters on whiskers of box plots in (**B**) and error bars in (**C**) & (**D**) indicate statistically significant differences (one-way ANOVA followed by post-hoc Tukey’s test, p <0.05). Scale bar=1cm in (**A**).

### LDL1 and LDL2 interact with FVE to regulate flowering time in *Arabidopsis*

^24, 25^. A recent report showed that LDL1 and LDL2 interact with HDA6 to regulate the circadian clock of *Arabidopsis* ^33^. HDA6 is a crucial part of the autonomous flowering pathway and is reported to act in a multiprotein complex that includes HDA5, FVE, and FLD as well. This multiprotein complex is established to induce flowering and the repression of FLC, MAF4, and MAF5 expression ^24, 25^. Therefore, we checked the expression of *MAF4* and *MAF5* in *ldl1ldl2hda6* triple mutant plants and their transcript levels were even more elevated than in *ldl1ldl2* double mutant plants. This indicates that LDL1 and LDL2 might be a part of a much bigger co-repressor complex that represses various MADS-box transcription factors to induce flowering. To further confirm this hypothesis, we checked the one-to-one interaction of LDL1 and LDL2 with each other and different known members of the co-repressor complex, FLD, HDA5, and FVE using yeast-2 hybrid (Y2H) assay (Figure 6A and Supplementary Figure5). We found that both LDL1 and LDL2 interact with FVE in the Y2H assay indicating the specificity of the interaction. To validate the interaction of LDL1 and LDL2 with FVE at a genetic level, we generated *fve*^*C*^ single mutant and *ldl1fve*^*C*^ and *ldl2fve*^*C*^ double mutant plants utilizing CRISPR-mediated genome editing. We selected the *fve*^*C*^, *ldl1fve*^*C*^, and *ldl2fve*^*C*^ T1 plants with a similar deletion in exon 1 of *FVE* for analyzing alteration in their flowering time (Supplementary Figure6). We found the *fve*^*C*^ single mutant plants showed late-flowering phenotype as compared to the WT plants (Figure 6C). *ldl2fve*^*C*^ double mutant plants flowered later than *fve*^*C*^ single mutant plants and *ldl1fve*^*C*^ double mutant plants showed delayed flowering even compared to *ldl2fve*^*C*^ plants (Figure 6D). Therefore, our results suggest that LDL1 and LDL2 are potential members of the corepressor complex through FVE and HDA6, and FVE promotes bolting co-dependently with LDL1 and LDL2.

**Figure 6.**
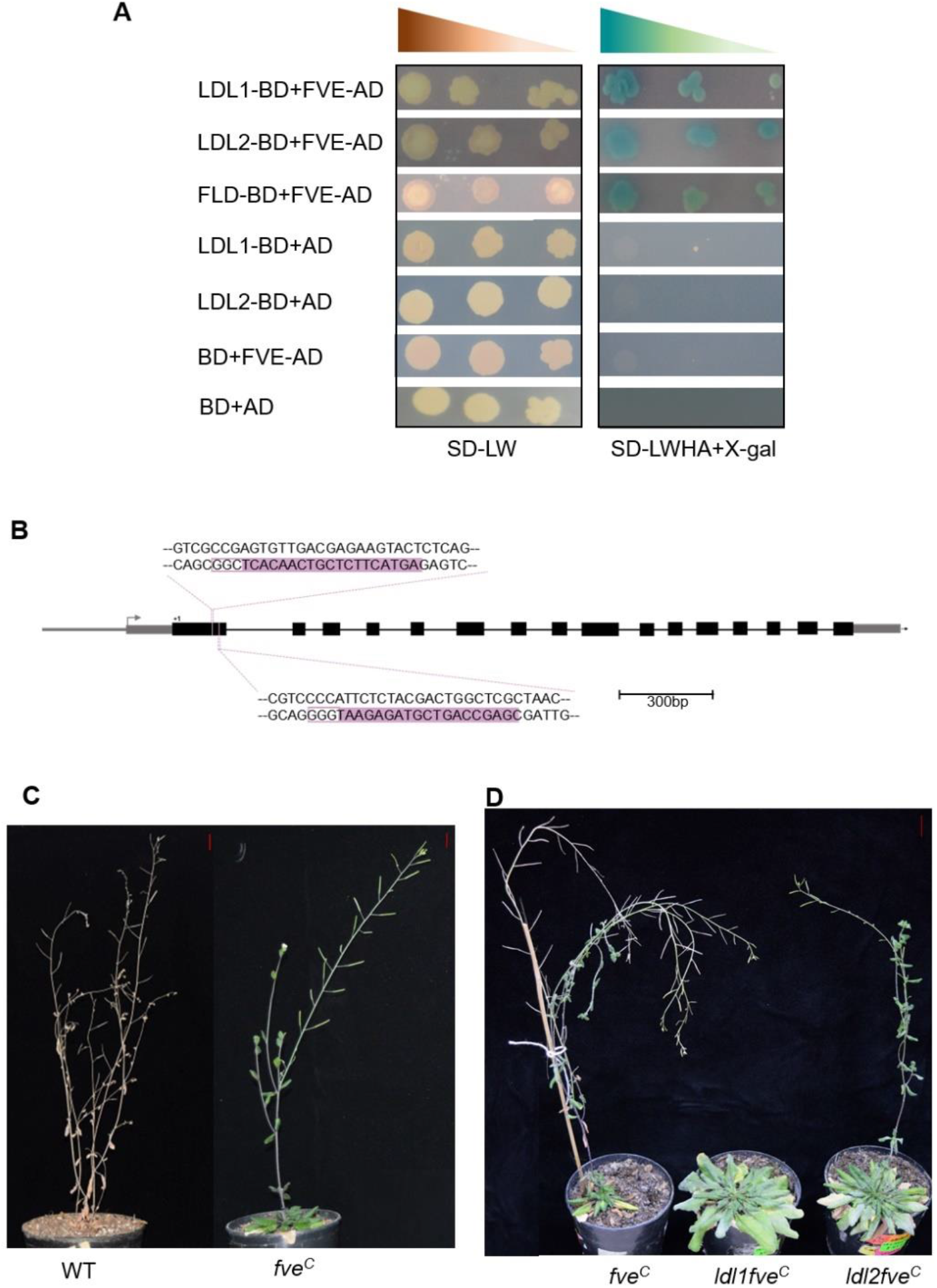
LDL1 and LDL2 interact with FVE to induce flowering. (**A**) LDL1 and LDL2 interact with FVE in a yeast two-hybrid assay. Co-transformed yeast cells were grown on SD medium lacking Leu and Trp (SD-LW) and the interaction was detected by the growth of yeast cells on quadruple dropout medium supplemented with 5-bromo-4-chloro-indolyl-galactopyranoside (SD-LWHA+X-gal). The blue colour indicates MEL1 protein activity. FLD-BD and FVE-AD were taken as positive controls and empty vectors BD and AD were taken as negative controls. The experiment was repeated three times. (**B**) Map showing the position of two guide RNAs in *FVE* gene for mutating FVE protein using CRISPR-Cas9. (**C**) Late flowering phenotype of *fve* with respect to the WT plant. (**D**) Late flowering phenotype of *ldl1fve* and *ldl2fve* with respect to *fve* plant. Scale bar=1cm in (**C**) and (**D**).

### MAF5 interacts with FLC and SVP to regulate the expression of FT

MADS-box genes are known to form multimeric protein complexes ^16^. FLC is known to interact with another MADS-box gene SVP to negatively regulate the expression of floral activators *FT* and *SOC1* ^7-9, 19^. Lately, MAF4 has been shown to interact with FLC and SVP and regulate the expression FT ^15^. In contrast, not much is known about MAF5 in terms of its interacting partners and direct downstream targets. Using the Y2H assay we found that MAF5 interacts with FLC and SVP and forms dimer with itself (Figure 7A). Since MAF5 is the clade member of FLC and interacts with both FLC and SVP, it is possible that MAF5 also binds to the chromatin of *FT* to repress the expression of FT and hence flowering. We quantified the promoter activity of *proFT:LUC* in *N*.*benthamiana* leaves using FLC, SVP, and MAF5 in different combinations.. FLC and SVP co-infiltrated together repressed *FT* promoter activity in contrast to FLC alone (Figure 7B). We also found that promoter activity of *FT* was reduced in the presence of MAF5 in combination with FLC and SVP as compared to FLC and SVP alone (Figure 7C and 7D). Our results suggest that the interaction of MAF5 with FLC and SVP represses *FT* in an additive manner to repress flowering.

**Figure 7.**
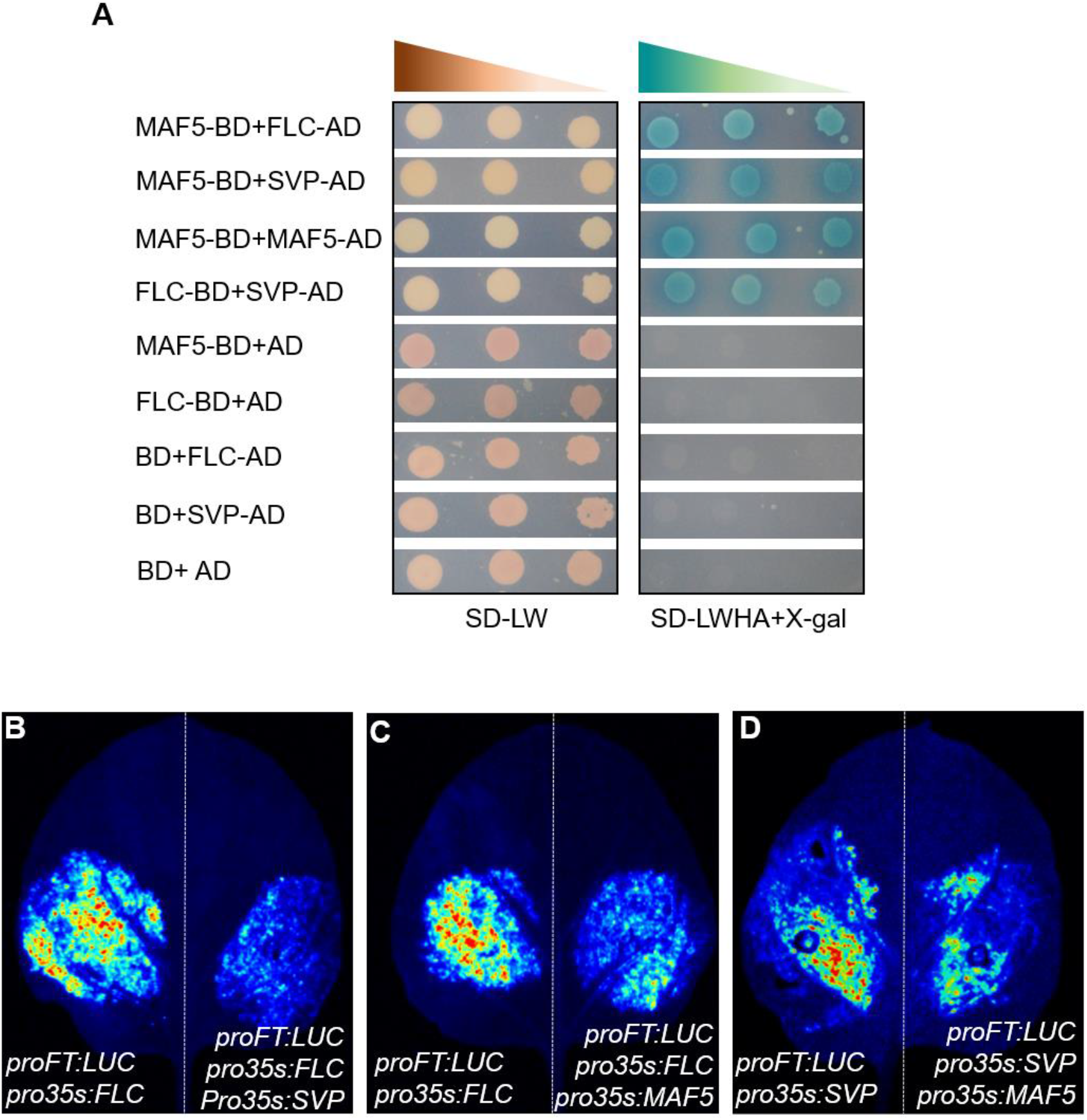
MAF5 interacts with FLC and SVP and negatively regulates *FT* expression. (**A**) MAF5 forms dimers with itself and interacts with FLC and SVP in a yeast two-hybrid assay. Co-transformed yeast cells were grown on SD-LW and the interaction was detected by the growth of yeast cells on SD-LWHA+X-gal medium. The blue colour indicates MEL1 protein activity. FLC-BD and SVP-AD were taken as positive controls and empty vectors BD and AD were taken as negative controls. The experiment was repeated three times. (**B, C & D**) Luciferase transactivation assay showing relative activity of *FT* promoter *in Nicotiana benthamiana* leaves co-infiltrated with different transcription factors. The experiment was repeated three times.

## Discussion

The seed of this study was planted by a genetic study, where we found that *ldl1flc* flowered significantly earlier than *ldl1* single mutant but later than the *flc* single mutant (Supplementary Figure 1). This observation implied the presence of additional targets of LDL1, which could contribute to its role in the regulation of flowering time. FLD, a family member of LDL1 is a well-known part of a corepressor complex, which functions in the autonomous flowering pathway and represses *FLC* and its clade members *MAF4* and *MAF5* to induce flowering ^24^. Expression of *FLC, MAF4*, and *MAF5* was upregulated in *ldl1* mutant and the expression of *FLC* and *MAF4* was downregulated in *LDL1 OE* plants indicating that LDL1 negatively regulates the expression of *FLC, MAF4*, and *MAF5* (Figure 1 A and 1B). We didn’t find any significant reduction in *MAF5* transcripts in *LDL1 OE* plants than in the WT plants (Figure 1B). The possible reason could be that *MAF5* is already highly repressed in the WT at 12 dag to induce the expression of floral integrator genes. There are several inconsistencies regarding the role of MAF4 and MAF5 as floral repressors. A primary study done by Ratcliffe et al. suggested that generating *MAF4* and *MAF5* overexpression in the Col-0 background either had no visible phenotype or flowered earlier than the Col-0 plants. However, generating their overexpression in the *ler* background significantly delayed the flowering time ^13^. Later it was found that T-DNA insertion mutant *maf4* in Col-0 background shows early flowering under short days ^14^. In contrast to its clade members, *FLC, MAF1*-*MAF3* which are downregulated when subjected to vernalization, *MAF4* and *MAF5* show somewhat dynamic expression patterns ^42^. To understand their role as floral repressors, we observed the flowering phenotype of *maf4* and *maf5* mutants in the Col-0 background under long-day conditions, and both *maf4* and *maf5* showed early flowering phenotype as compared to the WT plants proving their role as a potent floral repressor (Figure 1C to 1E). In winter annual ecotypes of *Arabidopsis*, the presence of a dominant allele of *FRIGIDA* (*FRI*) contributes to higher levels of *FLC* so that the plants can surpass the winters and flower in spring ^5, 6^. A recent study also showed that expressing *FRIGIDA* under root-specific promoter in *Arabidopsis* leads to the upregulation of *MAF4* and *MAF5* in the root, which might result in the formation of some mobile signal, which travels from root to shoot to antagonize the expression of *FT*, and hence delays flowering ^43^. This study also reinforces the role of MAF4 and MAF5 as floral repressors.

LDL1 is a histone modifier, and histone modifiers regulate the expression of their direct targets by binding to their chromatin and changing chromatin marks. Using *proLDL1:LDL1-GUS* plants, LDL1 was found to be enriched on the promoter and exon1 of *MAF4* and *MAF5* (Figure 2A and 2B). Genome-wide ChIP-seq analysis of LDL1 also revealed that 30 to 35% of LDL1 binding sites are present on the promoters and 30% to 40% are present in the first exon of protein-coding genes, which also aligns with our result ^44^. The presence of GUS activity of *proMAF4:GFP-GUS* and *proMAF5:GFP-GUS* in *ldl1* mutant plants (Figure 2D to 2G) and the rescued late-flowering phenotype of *ldl1* plants by the mutation in *MAF4* and *MAF5* loci (Figure 3 further confirm the repression of *MAF4* and *MAF5* by LDL1. LDL1 was found to have invitro H3K4me2 demethylase activity (Figure 4B and 4C) and thus it demethylates H3K4me2 marks to repress its targets. In addition to H3K4me2 demethylase activity, LDL1 also possesses H3K9me2 demethylase activity (Figure 4B and 4D). During *in vitro* enzymatic assays Human LSD1 was found only to demethylate only H3K4me1/me2 marks, but the interaction of LSD1 with certain partners resulted in demethylation of H3K9 *in vivo* ^39–41^. Interestingly we found that, unlike LSD1, LDL1 can demethylate H3K9me2 marks independent of any interacting partners, indicating that LDL1 could also contribute to transcriptional activation. Interestingly a study came out showing LDL1 positively regulates the expression of *ANGUSTIFOLIA3* (*AN3*) and H3K9 methylation marks were found to be reduced on *AN3* loci in the *LDL1 OE* plants ^45^. Hence, LDL1 can remove both transcriptionally permissive and repressive marks in *Arabidopsis*, but the role of LDL1 as a transcriptional activator needs further exploration.

Apart from LDL1, LDL2 also represses the expression of *MAF4* and *MAF5*, and they do so in a concerted manner (Figure5). When this manuscript was under preparation, both LDL1 and LDL2 were shown to interact with HDA6 to regulate circadian rhythm ^46^. The transcript level of both *MAF4* and *MAF5* was more upregulated in *ldl1ldl2hda6* than *ldl1ldl2*, which was comparable to their flowering phenotype (Figure 5). Since HDA6 is also a part of the autonomous flowering pathway, its interaction with LDL1 and LDL2 and cumulative effect on the expression of *MAF4* and *MAF5* and hence on floral transition proposes LDL1 and LDL2 as potential members of the autonomous flowering pathway. This hypothesis was confirmed by the direct one to one interaction of LDL1 and LDL2 with a crucial component of autonomous pathway, FVE (Figure 6A). *FVE* was identified as one of the first loci of the autonomous flowering pathway through genetic screening ^47^. It is a homolog of mammalian RETINOBLASTOMA-ASSOCIATED PROTEINS RBAP46 AND RBAP48 (RbR) and yeast MULTICOPY SUPPRESSOR OF IRA1(MSI), which are involved in chromatin modifications ^48^. These proteins contain several repeats of the WD40 motif, which allows protein-protein interaction. These proteins have no enzymatic activity but are involved in stabilizing various chromatin-modifying complexes (Summarised in supplementary table 1) The interaction of LDL1 with FVE was then confirmed in planta, by generating *fve*^*c*^, *ldl1fve*^*c*^, and *ldl2fve*^*c*^ plants. We observed that *fve*^*c*^ single mutant plants showed a late flowering phenotype (Figure 6C) as observed by Koornneef et al ^47^. *ldl2fve*^*c*^ double mutant plants flowered later than *fve*^*c*^ and the flowering in *ldl1fve*^*c*^ plants was even more delayed than *ldl2fve*^*c*^ plants, consistent with the flowering phenotype of *ldl1* and *ldl2* plants (Figure 6D and5A). In *Arabidopsis*, FVE interacts with HDA5, HDA6, and FLD to repress the expression of *FLC, MAF4*, and *MAF5* to induce flowering ^24, 25^. This suggests that LDL1 and LDL2 could be a part of the autonomous flowering pathway through their interaction with FVE and HDA6.

Unlike *FLC* and its clade members, *MAF1-MAF3* which are downregulated during vernalization, expression of *MAF4* and *MAF5* show a dynamic pattern during vernalization, where their expression first increases during vernalization and then decreases ^49^. Another study reported that *NAT-lncRNA_2962* (*MAS*), antisense long non-coding RNA (lncRNA) produced by the *MAF4* locus is induced by cold treatment and positively regulates the expression of *MAF4* ^50^. Therefore, it is possible that the expression of *MAF4* and *MAF5* is positively regulated by their respective lncRNAs under vernalization to avoid precocious flowering. Contrastingly, the lncRNAs produced by *FLC* under cold treatment, COOLAIR, COLDAIR, and COLDWRAP repress the expression of *FLC* ^51–53^. These opposite expression patterns of *FLC* and *MAF4* under vernalization are the outcome of differential regulation by their respective lncRNAs and thus highlights the importance of lncRNAs in regulating flowering time. Recently, a report from Hung et al. showed increased levels of lncRNAs produced by *MAF4* and *MAF5* loci independent of cold exposure in *ldl1ldl2hda6* plants ^44^. Combined, these observations direct to the possibility that the co-repressor complex involving LDL1 and LDL2, apart from regulating levels of MAF4 and MAF5 by changing their chromatin status directly, also regulates them indirectly through their corresponding lncRNAs.

MAF5 controls floral transition by interacting with FLC and SVP to negatively regulate the expression of *FT* (Figure 7). Recent advances have shown that SVP interacts with FLC, MAF2, and MAF4 to repress floral transition ^15, 19^. Our results indicate that MAF5 could also be a part of SVP-FLC-MAFs tetrameric complexes. Several other MADS-box transcription factors, AGAMOUS (AG), SEPLATTA3 (SEP3), APETALLA1(AP1), AP3, and FRUITFUL (FUL) act in tetrameric complexes to allow floral organ initiation ^54^. Therefore, it is possible that several MADS-box complexes coexist in the nucleus of a cell and compete for CArG motifs on their downstream genes and the abundance of the specific complex at a given time or their tissue specific expression would decide the fate of floral transition and floral organ development.

To summarise our work, we found that LDL1 promotes floral transition in *Arabidopsis* by suppressing the expression of floral repressors *MAF4* and *MAF5*. LDL1 has H3K4me2 demethylation activity and binds to *MAF4* and *MAF5* chromatin to alter their chromatin status. LDL2 also represses MAF4 and MAF5 expression, and both histone demethylases interact with FVE, and thus potentially act as parts of a bigger corepressor complex involved in the autonomous flowering pathway. MAF5 interacts with two other MADS-Box floral repressors, FLC and SVP and together they bind to the promoter of FT to inhibit its expression to hinder precocious flowering (Figure 8).

**Figure 8.**
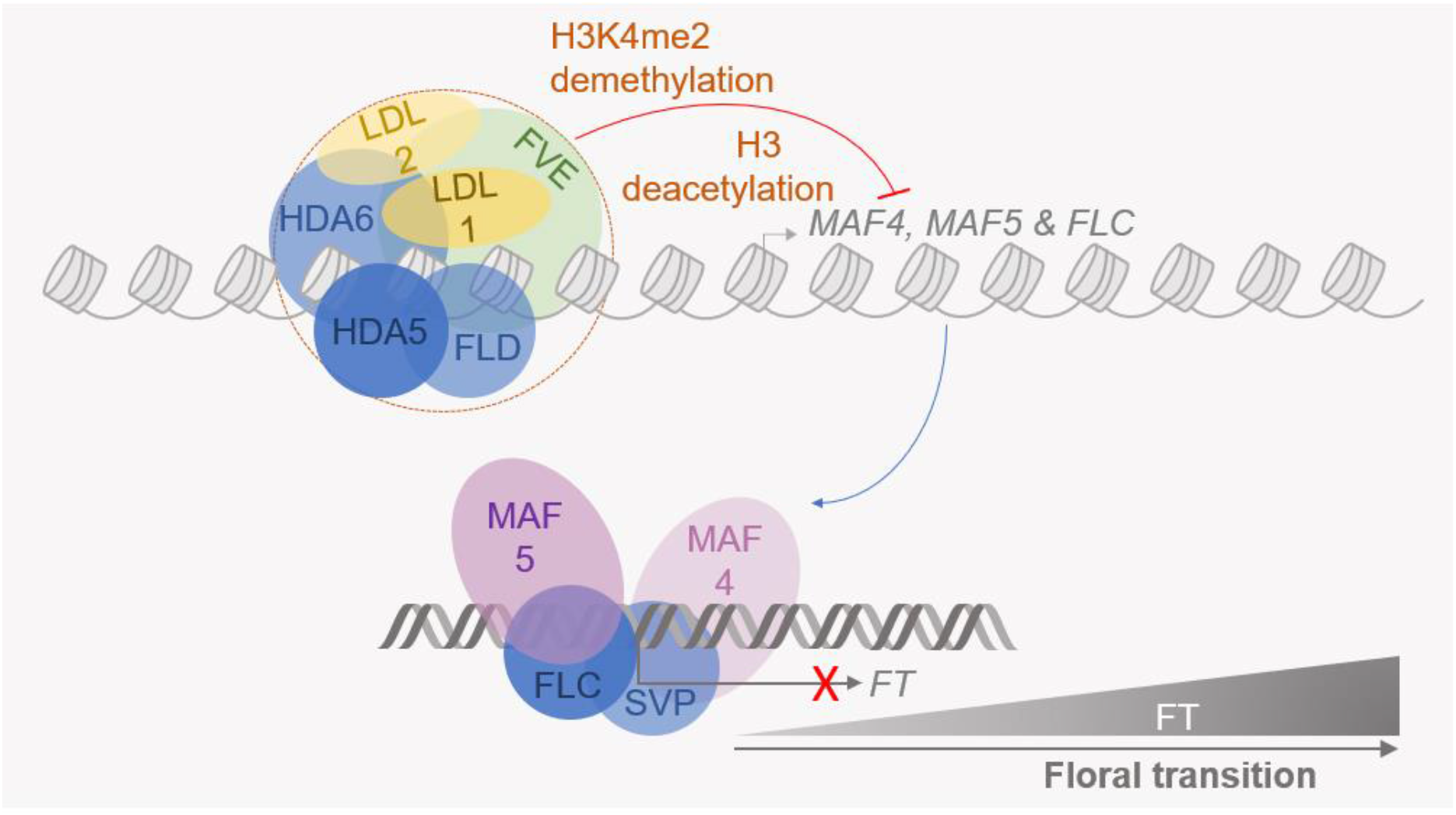
LDL1/LDL2 interact with FVE and promote floral transition mutually by repressing MADS-box transcription factors. LDL1 and LDL2 interact with FVE and HDA6 and assemble as a part of a corepressor complex on *MAF4, MAF5*, and *FLC* chromatin. This corepressor complex alters the chromatin state of these loci to suppress transcription. MAF4 and MAF5 further interact with other MADS-box transcription factors, like FLC and SVP. These combinations of MADS-box proteins bring down the transcriptional output of the *FT* locus, and thus, affect vegetative to floral transition.

## Materials and Methods

### Plant material and growth conditions

Arabidopsis thaliana Wild type Columbia-0 (WT), *ldl1-1* (SALK_142477), *ldl1-2* (SALK_034869C), *flc-3, maf4* (SAIL_1213_A08), *maf5* (SALK_015513), *ldl2* (SALK_135831C), *ldl1flc, ldl1maf4, ldl1maf5, ldl1ldl2* and *ldl1ldl2hda6* were used in the study. *Arabidopsis thaliana* seeds were surface sterilized with 70 % ethanol containing 0.1 % (v/v) Triton X-100 in a microcentrifuge tube for 10 min followed by 5-6 times washes with sterile water. Surface sterilized seeds were kept at 4°C (in dark) for 3 days to synchronize germination. Seeds were then transferred on 0.5X Murashige and Skoog medium ^55^ plates containing 0.8 % (v/v) plant agar. Plates were kept in a near-vertical position in the plant growth chamber having 21°C temperature and light illumination (around 120 μM^−2^) period for 16 hrs followed by an 8hrs dark period.

For analyzing flowering phenotype plants were either grown on 0.5X Murashige and Skoog medium for 6 days and then transferred to pots or seeds were directly sprinkled in the pots, stratified for 3 days, and then transferred to the closed growth chamber. Rosette leaves were scored at the appearance of inflorescence. All the phenotypic experiments were repeated thrice.

### Transgenics generation

*LDL1 OE* plants were generated by amplifying 2535 bp of the coding sequence in modified pCAMBIA1301 vector under *pRPS5A* promoter. The construct was transformed into WT plants through the *Agrobacterium tumefaciens* (*GV3850*) mediated floral dip method ^56^. For constructing *proLDL1:LDL1-GUS* plants, we amplified 3357 bp of gDNA and cloned it in pCAMBIA1301. The construct and the empty vector control were transformed in *ldl1* plants. For generating the *proMAF4:GFP-GUS* construct, 1242bp upstream and 187 bp downstream of translation start site was taken in frame with GFP and for *proMAF5: GFP-GUS* construct, 1977bp upstream and 62 bp downstream of translation start site were taken in frame with GFP (pCAMBIA1304). Both *proMAF4:GFP-GUS* and *proMAF5: GFP-GUS* were transformed into WT and *ldl1* mutant backgrounds. For generating *fve* mutant plants using genome editing, we employed the system, which allowed the assembly of two guide RNAs (gRNAs) to maximize the probability of generating the mutant. We used http://www.rgenome.net/cas-offinder/ to evaluate target specificities to rule out the possibility of potential off-target and selected the gRNA with no or minimum off-targets. Both selected gRNAs targeted exon 1 of *FVE*. Using the golden gate assembly method, we cloned the two gRNAs in the binary vector and confirmed the clones using colony PCR and sequencing (Figure 6.7). The confirmed clone was transformed into WT, *ldl1*, and *ldl2* plants.

Positive plants were selected by growing the T1 seeds on Hygromycin B selection media. Resistant plants were grown and used for expression level analysis and histochemical GUS assay. Genetic segregation analysis was performed to confirm single T-DNA insertion and homozygous T3 seeds were used for further experiments.

### Histochemical GUS assay

To study the spatiotemporal expression of the various genes, we have performed GUS histochemical analysis as described previously ^57^. The *Arabidopsis* whole seedlings or other tissues were transferred in the microcentrifuge tubes containing an appropriate amount of GUS staining buffer [50 mM sodium phosphate (pH 7), 50 mM EDTA (pH 8), 0.5 mM K3Fe(CN)6, 0.1 % Triton X-100, 1 mM X-Gluc] and kept at 37°C and checked at regular intervals for the development of blue-colored end product as GUS enzyme cleaves the substrate, X-Gluc. Once an adequate signal had developed in the different tissues under study, the GUS staining buffer was replaced with a solution of acetone: ethanol (1:3 ratio) to remove chlorophyll from the green tissues. Desired tissues were placed on slides having diluted chloral hydrate solution and images were taken with the help of a stereo-zoom microscope (Nikon AZ100, Tokyo, Japan).

### Gene expression analysis

Expression of all flowering-related genes was checked in the shoot of 12 days old seedlings. RNA was isolated using Trizol (Sigma) as per the manufacturer’s guidelines. RNA was reverse transcribed using M-MLV RT (Thermo Scientific) and Real-Time Quantitative Reverse Transcription PCR (qRT PCR) was performed in the “7900HT FAST” real-time PCR system (Applied Biosystems) using SYBR green based assay. UBIQUITIN5 (UBQ5), and ACTIN7 (ACT7) were used as endogenous controls. The sequences of primers used for qRT-PCR are provided in **Table S2**.

### Chromatin immunoprecipitation (ChIP) and qRT PCR

The ChIP was performed as described ^58^(). 2g of sample was harvested and fixed in a formaldehyde-based buffer. Chromatin was sheared to an average length of 500 bp and immunoprecipitated with specific antibodies, anti-GUS, and IgG. Immunoprecipitated chromatin was quantified using qRT-PCR and normalized with respect to *ACTIN7*.

### Detection of LDL1 demethylase activity

LDL1 CDS was cloned in the pGEX-4T1 vector (GST tag). Purified LDL1 protein was incubated with calf thymus histones (Sigma) at RT for 4h in the presence of 30 % glycerol and 50mM Tris-HCl (pH 8) at room temperature (RT) for 4 h. Histone demethylase activity of LDL1 was then evaluated by western blot using H3K4me1, H3K4me2, H3K9me1, and H3K9me2 specific antibodies (Abcam) as per Abcam manual. We used alkaline phosphatase (Sigma) as a secondary antibody. Detection was done using 5-Bromo-4-chloro-3-indolyl phosphate/ nitro blue tetrazolium (BCIP/NBT) solution. BCIP is a substrate of alkaline phosphatase and catalyzed BCIP reacts with NBT to produce a dark blue insoluble precipitate. To stop the reaction, the blot was washed with autoclaved water.

### Yeast two-hybrid (Y2H) analysis

The coding sequence of LDL1, LDL2, FLD, FVE, MAF5, FLC, and SVP was cloned in gateway-based pGBKT7g and pGADT7g Y2H vectors. Positive clones were transformed in *Saccharomyces cerevisiae* strain Y2H gold cells (Takara biotech) and plated on SD-LEU-TRP (DDO) plates. Yeast transformation was performed as per the manufacturer’s protocol (EZ-Yeast transformation kit, MP Biomedical, USA). Colonies obtained on DDO plates were streaked on SD–ADE-HIS-LEU-TRP medium plates containing X-α-gal (QDOX) plates. Plates were incubated at 30 °C for 3-5 days.

### Luciferase assay

For generating *proFT:LUC*, 1688 bp upstream of the translation start site was amplified and cloned in pGREENII0800. Coding sequences of MAF5, SVP, and FLC were cloned under CaMV 35S promoter in pGWB441. Constructs were transformed into Agrobacterium tumefaciens (GV3101). Constructs were coinfilterated into *Nicotiana benthamiana* with different combinations and luciferase activity of *proFT:LUC* was detected using its substrate luciferin after 2 days using chemidoc (BioRad).

### Statistical Analysis

Numerical data from all experiments were represented with Microsoft Excel. Student’s t-tests were done using Microsoft Excel and One-way ANOVA and post hoc Tukey’s tests were done using IBM SPSS software. Details of the error bar, replicates, statistical tests applied, and significances are mentioned in the relevant figure legends.

### Accession numbers

LDL1 (AT1G62830), LDL2 (AT3G13682), FLD (AT3G10390), FVE (AT2G19520), HDA5 (AT5G61060), HDA6 (AT5G63110), FLC (AT5G10140), SVP (AT2G22540), MAF4 (AT5G65070) and MAF5 (AT5G65080)

## Acknowledgements

We acknowledge NIPGR for providing internal funding and the Central Instrument Facility of NIPGR to carry out this work. We acknowledge the DBT-eLibrary Consortium (DeLCON) for providing access to e-resources. Ma, SC, and SS acknowledge the Council of Industrial and Scientific Research (CSIR; Govt. of India) for providing fellowships. AKS acknowledges funding from the Science and Engineering Research Board (SERB), Department of Science and Technology (DST), Government of India (research grant no. EMR/2016/002438). We thank Prof. Richard Amasino for *fld-1flc-3* and Dr. Keqiang Wu for *ldl1ldl2* and *ldl1ldl2hda6* seeds.

## Author contributions

AKS conceived the original research plan and design, supervised the work, and revised the manuscript. Ma designed and performed most of the experiments, analyzed data, and wrote the manuscript. SC and SS made substantial contributions to the experiments and complemented the writing of the manuscript.

## Conflict of interest

All authors have read and approved the manuscript and declare no conflict of interest.

Supplementary Figure 1. ***ldl1flc* double mutant flowers earlier than *ldl1* single mutant but later than *flc* single mutant**. (**A**) Flowering phenotype of *ldl1flc* with respect to *ldl1* and *flc*. (**B**) Rosette leaf numbers of *ldl1, ldl1flc*, and *flc* plants at bolting (n=15). (**C**) Days taken to flower by *ldl1, ldl1flc*, and *flc* plants (n=15). Different letters on whiskers of box plots in (**B**) and (**C**) indicate statistically significant differences (one-way ANOVA followed by post-hoc Tukey’s test, p <0.05). Scale bar=1cm in (**A**).

Supplementary Figure 2. **Development of *LDL1 Oe* construct and transgenic lines**. (**A**) Construct map for *LDL1* overexpression. (**B**) *LDL1Oe#1 and #4* showed maximum overexpression, respectively. Error bars indicate the standard deviation (± SD) of three technical replicates.

Supplementary Figure 3. **Translational fusion (*proLDL1:LDL1-GUS*) complements late flowering phenotype of *ldl1***. (**A**) *LDL1* expression level in translational fusion line of LDL1 and empty vector (EV) control in *ldl1* background. (B) Rosette leaf numbers of *EV* (*ldl1*) and *proLDL1-GUS* (*ldl1*) plants at bolting (n=15). The experiment was repeated thrice. Error bars indicate the standard error (± SE) of three independent experiments. Asterisks indicate significant differences (\**p* ≤ 0.05, \*\**p* ≤ 0.01, \*\*\**p* ≤ 0.001; unpaired two tailed *t-*test). Scale bar=1cm in (**C**).

Supplementary Figure 4. **Tissue specific expression pattern of LDL1**. (**A**) Construct showing translational fusion map of *LDL1*. **(B)** GUS activity was observed in different parts of *pLDL1:LDL1-GUS (ldl1)* transgenic line. (i) SAM of four days old seedling (arrowhead indicates SAM) (ii) young leaves, (iii) flowers, (iv) primary root, (v) stage iv LRP, (vi) stage vii LRP, (vii) LR. Scale bar 2 mm in (i) and (iii) and 50 μm in (ii), (iv) – (vii).

Supplementary Figure 5. **LDL1 doesn’t interacts with LDL2, FLD and HDA5 in Y2H assay**. Co-transformed yeast cells were grown on SD-LW and the interaction was detected by the growth of yeast cells on SD-LWHA+X-gal medium and the blue colour indicating MEL1 protein activity. p53-BD and SV40-AD were taken as positive controls and empty vectors were taken as negative controls.

Supplementary Figure 6. **CRISPR/Cas9 mediated mutagenesis in *FVE*** (**A**) Vector map of CRISPR construct. (**B**) Sequencing alignment of gRNA target region showing deletion in *FVE* in the WT, *ldl1* and *ldl2* background.

Supplementary Table 1. Summary of FVE homologs as a part of different chromatin modifying complexes in *Arabidopsis thaliana* (*At*), *Saccharomyces cerevisiae* (*Sc*), *Drosophila melanogaster* (*Ds*) and *Homo Sapiens* (*Hs*)

Supplementary Table 2. List of primers used in the study

## References

1. Franks, S.J. The unique and multifaceted importance of the timing of flowering. American journal of botany 102, 1401–1402 (2015).

2. He, Y. & Amasino, R.M. Role of chromatin modification in flowering-time control. Trends in plant science 10, 30–35 (2005).

3. Srikanth, A. & Schmid, M. Regulation of flowering time: all roads lead to Rome. Cellular and molecular life sciences 68, 2013–2037 (2011).

4. Zhang, Y., Zhou, Y., Chen, Q., Huang, X. & Tian, C.e.J.C.B.o.B. Molecular basis of flowering time regulation in Arabidopsis. 49, 469 (2014).

5. Michaels, S.D. & Amasino, R.M. FLOWERING LOCUS C encodes a novel MADS domain protein that acts as a repressor of flowering. Plant Cell 11, 949–956 (1999).

6. Sheldon, C.C. et al. The FLF MADS box gene: a repressor of flowering in Arabidopsis regulated by vernalization and methylation. Plant Cell 11, 445–458 (1999).

7. Helliwell, C.A., Wood, C.C., Robertson, M., James Peacock, W. & Dennis, E.S. The Arabidopsis FLC protein interacts directly in vivo with SOC1 and FT chromatin and is part of a high-molecular-weight protein complex. The Plant Journal 46, 183–192 (2006).

8. Searle, I. et al. The transcription factor FLC confers a flowering response to vernalization by repressing meristem competence and systemic signaling in Arabidopsis. Genes Dev 20, 898–912 (2006).

9. Li, D. et al. A repressor complex governs the integration of flowering signals in Arabidopsis. Dev Cell 15, 110–120 (2008).

10. Ratcliffe, O.J., Nadzan, G.C., Reuber, T.L. & Riechmann, J.L. Regulation of flowering in Arabidopsis by an FLC homologue. Plant Physiol 126, 122–132 (2001).

11. Scortecci, K., Michaels, S.D. & Amasino, R.M. Genetic interactions between FLM and other flowering-time genes in Arabidopsis thaliana. Plant Molecular Biology 52, 915–922 (2003).

12. Lee, J.H. et al. Regulation of temperature-responsive flowering by MADS-box transcription factor repressors. Science 342, 628–632 (2013).

13. Ratcliffe, O.J., Kumimoto, R.W., Wong, B.J. & Riechmann, J.L. Analysis of the Arabidopsis MADS AFFECTING FLOWERING gene family: MAF2 prevents vernalization by short periods of cold. The Plant cell 15, 1159–1169 (2003).

14. Gu, X., Jiang, D., Wang, Y., Bachmair, A. & He, Y. Repression of the floral transition via histone H2B monoubiquitination. The Plant journal : for cell and molecular biology 57, 522–533 (2009).

15. Gu, X. et al. Arabidopsis FLC clade members form flowering-repressor complexes coordinating responses to endogenous and environmental cues. Nature Communications 4, 1947 (2013).

16. de Folter, S. et al. Comprehensive interaction map of the Arabidopsis MADS Box transcription factors. Plant Cell 17, 1424–1433 (2005).

17. Riechmann, J.L., Krizek, B.A. & Meyerowitz, E.M. Dimerization specificity of Arabidopsis MADS domain homeotic proteins APETALA1, APETALA3, PISTILLATA, and AGAMOUS. Proceedings of the National Academy of Sciences of the United States of America 93, 4793–4798 (1996).

18. A Hepworth, S.R., Valverde, F. Ravenscroft D., Mouradov, A. & Coupland, G. Antagonistic regulation of flowering-time gene SOC1 by CONSTANS and FLC via separate promoter motifs. Embo j 21, 4327–4337 (2002).

19. Lee, J.H. et al. Role of SVP in the control of flowering time by ambient temperature in Arabidopsis. Genes Dev 21, 397–402 (2007).

20. Whittaker, C. & Dean, C. The FLC Locus: A Platform for Discoveries in Epigenetics and Adaptation. 33, 555–575 (2017).

21. Cheng, J.Z., Zhou, Y.P., Lv, T.X., Xie, C.P. & Tian, C.E. Research progress on the autonomous flowering time pathway in Arabidopsis. Physiol Mol Biol Plants 23, 477–485 (2017).

22. He, Y., Michaels, S.D. & Amasino, R.M. Regulation of Flowering Time by Histone Acetylation in <em>Arabidopsis</em>. Science 302, 1751–1754 (2003).

23. Domagalska, M.A. et al. Attenuation of brassinosteroid signaling enhances <em>FLC</em> expression and delays flowering. Development 134, 2841–2850 (2007).

24. Yu, C.-W. et al. HISTONE DEACETYLASE6 Interacts with FLOWERING LOCUS D and Regulates Flowering in Arabidopsis. Plant Physiology 156, 173–184 (2011).

25. Luo, M. et al. Regulation of flowering time by the histone deacetylase HDA 5 in A rabidopsis. The Plant Journal 82, 925–936 (2015).

26. Gu, X. et al. Arabidopsis homologs of retinoblastoma-associated protein 46/48 associate with a histone deacetylase to act redundantly in chromatin silencing. PLoS Genet 7, e1002366 (2011).

27. Jiang, D., Yang, W., He, Y. & Amasino, R.M. Arabidopsis relatives of the human lysine-specific Demethylase1 repress the expression of FWA and FLOWERING LOCUS C and thus promote the floral transition. The Plant Cell 19, 2975–2987 (2007).

28. Stavropoulos, P., Blobel, G. & Hoelz, A. Crystal structure and mechanism of human lysine-specific demethylase-1. Nature structural & molecular biology 13, 626–632 (2006).

29. Yang, M. et al. Structural basis for CoREST-dependent demethylation of nucleosomes by the human LSD1 histone demethylase. Molecular cell 23, 377–387 (2006).

30. Binda, C., Mattevi, A. & Edmondson, D.E. Structure-function relationships in flavoenzyme-dependent amine oxidations a comparison of polyamine oxidase and monoamine oxidase. Journal of biological chemistry 277, 23973–23976 (2002).

31. Binda, C., Newton-Vinson, P., Hubálek, F., Edmondson, D.E. & Mattevi, A. Structure of human monoamine oxidase B, a drug target for the treatment of neurological disorders. Nature structural biology 9, 22–26 (2002).

32. Zhao, M., Yang, S., Liu, X. & Wu, K. Arabidopsis histone demethylases LDL1 and LDL2 control primary seed dormancy by regulating DELAY OF GERMINATION 1 and ABA signaling-related genes. Frontiers in plant science 6, 159 (2015).

33. Hung, F.-Y. et al. The Arabidopsis LDL1/2-HDA6 histone modification complex is functionally associated with CCA1/LHY in regulation of circadian clock genes. Nucleic acids research 46, 10669–10681 (2018).

34. Krichevsky, A., Zaltsman, A., Kozlovsky, S.V., Tian, G.-W. & Citovsky, V. Regulation of root elongation by histone acetylation in Arabidopsis. Journal of molecular biology 385, 45–50 (2009).

35. Singh, S., Singh, A., Roy, S. & Sarkar, A.K. SWP1 negatively regulates lateral root initiation and elongation in Arabidopsis. Plant signaling & behavior 7, 1522–1525 (2012).

36. Krichevsky, A. et al. C2H2 zinc finger-SET histone methyltransferase is a plant-specific chromatin modifier. Developmental biology 303, 259–269 (2007).

37. Maruyama, D. et al. Independent control by each female gamete prevents the attraction of multiple pollen tubes. Dev Cell 25, 317–323 (2013).

38. Maiques-Diaz, A. & Somervaille, T.C.J.E. LSD1: biologic roles and therapeutic targeting. 8, 1103–1116 (2016).

39. Shi, Y. et al. Histone demethylation mediated by the nuclear amine oxidase homolog LSD1. Cell 119, 941–953 (2004).

40. Metzger, E. et al. LSD1 demethylates repressive histone marks to promote androgen-receptor-dependent transcription. Nature 437, 436–439 (2005).

41. Carnesecchi, J. et al. ERRα induces H3K9 demethylation by LSD1 to promote cell invasion. Proceedings of the National Academy of Sciences of the United States of America 114, 3909–3914 (2017).

42. Kim, D.-H. & Sung, S. Coordination of the Vernalization Response through a <em>VIN3</em> and <em>FLC</em> Gene Family Regulatory Network in <em>Arabidopsis</em>. The Plant Cell 25, 454–469 (2013).

43. Kong, X. et al. Expression of FRIGIDA in root inhibits flowering in Arabidopsis thaliana. Journal of Experimental Botany 70, 5101–5114 (2019).

44. Hung, F.-Y. et al. The expression of long non-coding RNAs is associated with H3Ac and H3K4me2 changes regulated by the HDA6-LDL1/2 histone modification complex in Arabidopsis. NAR Genomics and Bioinformatics 2 (2020).

45. Keren, I., Lapidot, M. & Citovsky, V. Coordinate activation of a target gene by KDM1C histone demethylase and OTLD1 histone deubiquitinase in Arabidopsis. Epigenetics 14, 602–610 (2019).

46. Hung, F.-Y. et al. The LDL1/2-HDA6 histone modification complex interacts with TOC1 and regulates the core circadian clock components in Arabidopsis. Frontiers in plant science 10, 233 (2019).

47. Koornneef, M., Hanhart, C. & Van der Veen, J. A genetic and physiological analysis of late flowering mutants in Arabidopsis thaliana. Molecular and General Genetics MGG 229, 57–66 (1991).

48. Kenzior, A.L. & Folk, W.R. AtMSI4 and RbAp48 WD-40 repeat proteins bind metal ions. FEBS letters 440, 425–429 (1998).

49. Kim, D.H. & Sung, S. Coordination of the vernalization response through a VIN3 and FLC gene family regulatory network in Arabidopsis. Plant Cell 25, 454–469 (2013).

50. Zhao, X. et al. Global identification of Arabidopsis lncRNAs reveals the regulation of MAF4 by a natural antisense RNA. Nat Commun 9, 5056 (2018).

51. Heo, J.B. & Sung, S. Vernalization-mediated epigenetic silencing by a long intronic noncoding RNA. Science 331, 76–79 (2011).

52. Kim, D.H. & Sung, S. Vernalization-Triggered Intragenic Chromatin Loop Formation by Long Noncoding RNAs. Dev Cell 40, 302–312.e304 (2017).

53. Helliwell, C.A., Robertson, M., Finnegan, E.J., Buzas, D.M. & Dennis, E.S. Vernalization-repression of Arabidopsis FLC requires promoter sequences but not antisense transcripts. PloS one 6, e21513–e21513 (2011).

54. Smaczniak, C. et al. Characterization of MADS-domain transcription factor complexes in Arabidopsis flower development. Proceedings of the National Academy of Sciences of the United States of America 109, 1560–1565 (2012).

55. A Classic Murashige, T. & Skoog, F. A revised medium for rapid growth and bioassays with tobacco tissue cultures. Physiologia Plantarum 15, 473–497 (1962).

56. Clough, S.J. & Bent, A.F. Floral dip: a simplified method for Agrobacterium-mediated transformation of Arabidopsis thaliana. The Plant journal : for cell and molecular biology 16, 735–743 (1998).

57. Jefferson, R.A., Kavanagh, T.A. & Bevan, M.W. GUS fusions: beta-glucuronidase as a sensitive and versatile gene fusion marker in higher plants. The EMBO journal 6, 3901–3907 (1987).

58. Saleh, A., Alvarez-Venegas, R. & Avramova, Z. An efficient chromatin immunoprecipitation (ChIP) protocol for studying histone modifications in Arabidopsis plants. Nature Protocols 3, 1018–1025 (2008).

59. Kaya, H. et al. FASCIATA genes for chromatin assembly factor-1 in arabidopsis maintain the cellular organization of apical meristems. Cell 104, 131–142 (2001).

60. Derkacheva, M. et al. Arabidopsis MSI1 connects LHP1 to PRC2 complexes. Embo j 32, 2073–2085 (2013).

61. Xu, Y. et al. MSI1 and HDA6 function interdependently to control flowering time via chromatin modifications. 109, 831–843 (2022).

62. Tan, L.-M. et al. Dual Recognition of H3K4me3 and DNA by the ISWI Component ARID5 Regulates the Floral Transition in Arabidopsis. The Plant Cell 32, 2178–2195 (2020).

63. Doyle, M.R. & Amasino, R.M. A Single Amino Acid Change in the Enhancer of Zeste Ortholog CURLY LEAF Results in Vernalization-Independent, Rapid Flowering in Arabidopsis. Plant Physiology 151, 1688–1697 (2009).

64. Yu, C.-W., Chang, K.-Y. & Wu, K. Genome-wide analysis of gene regulatory networks of the FVE-HDA6-FLD complex in Arabidopsis. Frontiers in plant science 7, 555 (2016).

65. Kaufman, P.D., Kobayashi, R. & Stillman, B. Ultraviolet radiation sensitivity and reduction of telomeric silencing in Saccharomyces cerevisiae cells lacking chromatin assembly factor-I. Genes Dev 11, 345–357 (1997).

66. Ge, Z., Wang, H. & Parthun, M.R. Nuclear Hat1p complex (NuB4) components participate in DNA repair-linked chromatin reassembly. The Journal of biological chemistry 286, 16790–16799 (2011).

67. Tie, F., Furuyama, T., Prasad-Sinha, J., Jane, E. & Harte, P.J. The Drosophila Polycomb Group proteins ESC and E(Z) are present in a complex containing the histone-binding protein p55 and the histone deacetylase RPD3. Development 128, 275–286 (2001).

68. Suganuma, T., Pattenden, S.G. & Workman, J.L. Diverse functions of WD40 repeat proteins in histone recognition. Genes Dev 22, 1265–1268 (2008).

69. Hassig, C.A., Fleischer, T.C., Billin, A.N., Schreiber, S.L. & Ayer, D.E. Histone deacetylase activity is required for full transcriptional repression by mSin3A. Cell 89, 341–347 (1997).

70. Sugimoto, N. et al. Cdt1-binding protein GRWD1 is a novel histone-binding protein that facilitates MCM loading through its influence on chromatin architecture. Nucleic Acids Res 43, 5898–5911 (2015).

71. Xue, Y. et al. NURD, a novel complex with both ATP-dependent chromatin-remodeling and histone deacetylase activities. Mol Cell 2, 851–861 (1998).

72. Kuzmichev, A., Nishioka, K., Erdjument-Bromage, H., Tempst, P. & Reinberg, D. Histone methyltransferase activity associated with a human multiprotein complex containing the Enhancer of Zeste protein. Genes Dev 16, 2893–2905 (2002).

73. Murzina, N.V. et al. Structural basis for the recognition of histone H4 by the histone-chaperone RbAp46. Structure (London, England : 1993) 16, 1077–1085 (2008).

74. Verreault, A., Kaufman, P.D., Kobayashi, R. & Stillman, B. Nucleosome assembly by a complex of CAF-1 and acetylated histones H3/H4. Cell 87, 95–104 (1996).

