## Supplementary data for "LDL1 and LDL2 histone demethylases interact with FVE to regulate flowering in *Arabidopsis*"

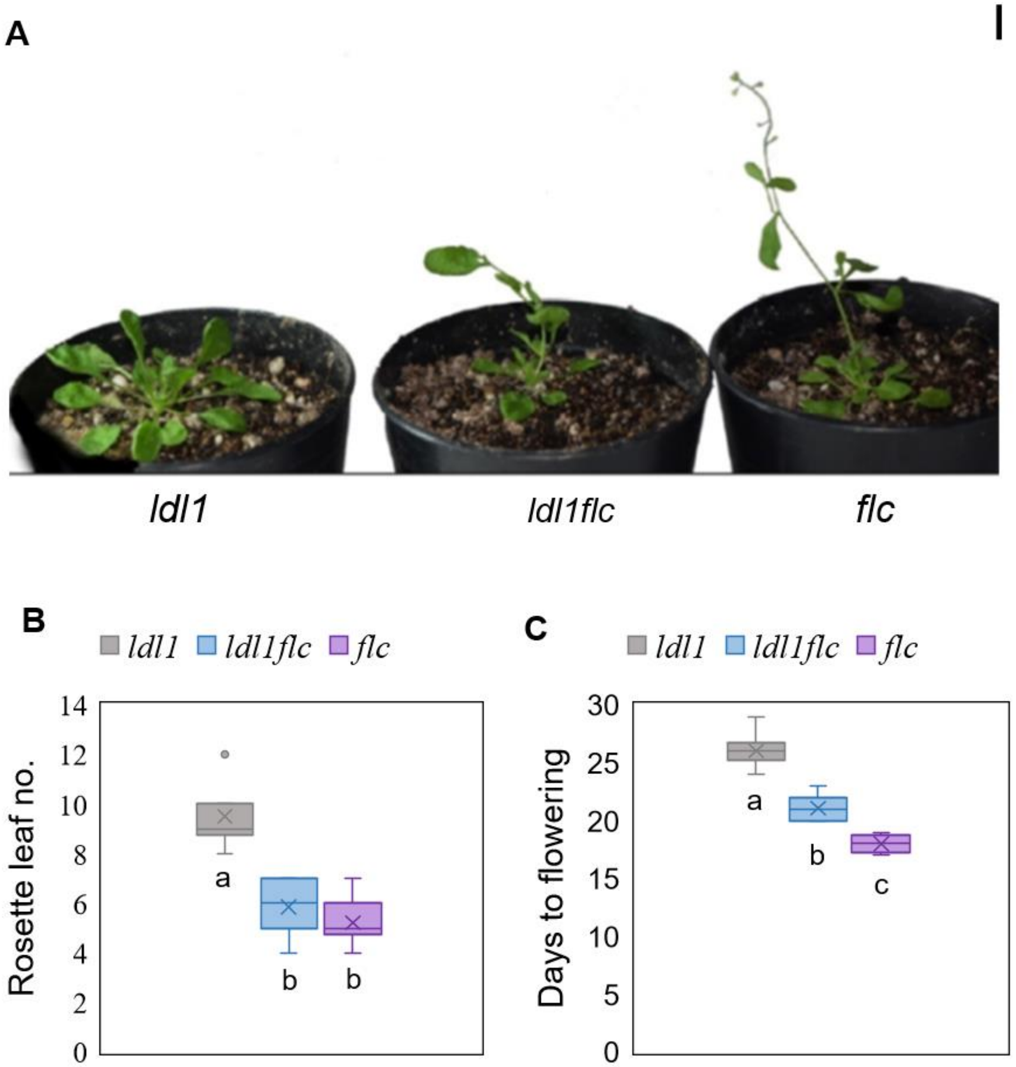


Supplementary Figure 1. ***ldl1flc* double mutant flowers earlier than *ldl1* single mutant but later than *flc* single mutant.** (**A**) Flowering phenotype of *ldl1flc* with respect to *ldl1* and *flc*. (**B**) Rosette leaf numbers of *ldl1*, *ldl1flc*, and *flc* plants at bolting (n=15). (**C**) Days taken to flower by *ldl1*, *ldl1flc*, and *flc* plants (n=15). Different letters on whiskers of box plots in (**B**) and (**C**) indicate statistically significant differences (one-way ANOVA followed by post-hoc Tukey's test, $p < 0.05$). Scale bar=1cm in (**A**).

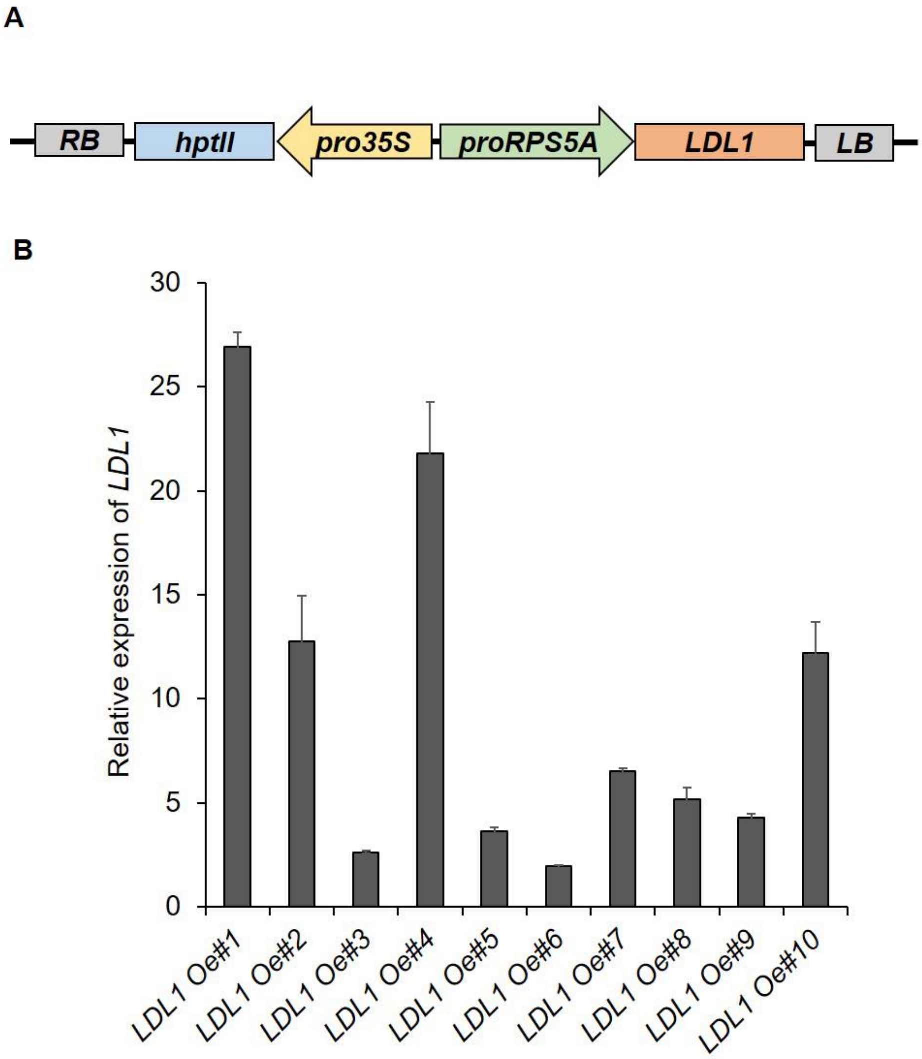


Supplementary Figure 2. **Development of *LDL1 Oe* construct and transgenic lines.** (**A**) Construct map for *LDL1* overexpression. (**B**) *LDL1Oe#1 and #4* showed maximum overexpression, respectively. Error bars indicate the standard deviation (± SD) of three technical replicates.

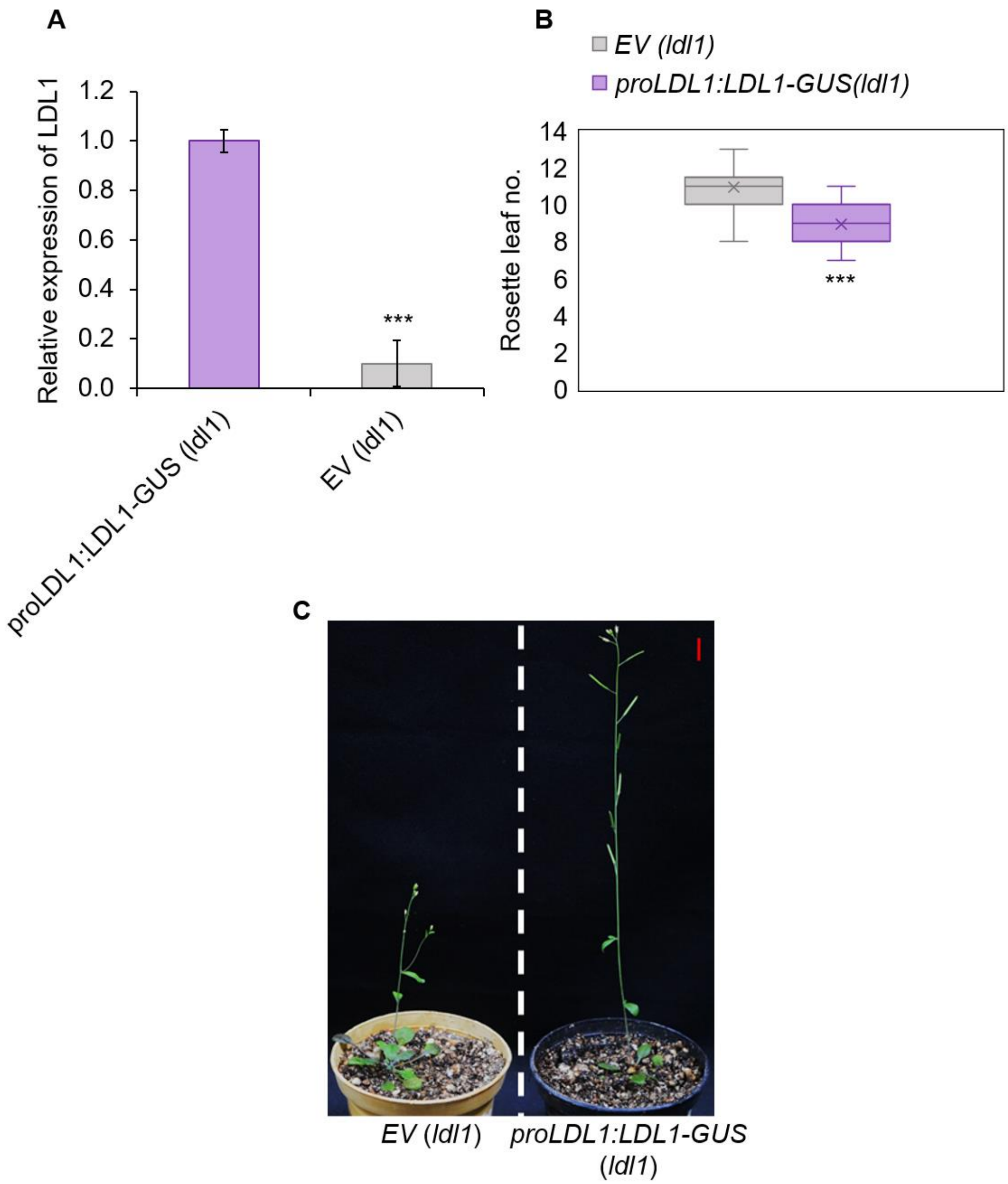


Supplementary Figure 3. **Translational fusion (*proLDL1:LDL1-GUS*) complements late flowering phenotype of *ldl1*.** (**A**) *LDL1* expression level in translational fusion line of LDL1 and empty vector (EV) control in *ldl1* background. (B) Rosette leaf numbers of *EV* (*ldl1*) and *proLDL1-GUS* (*ldl1*) plants at bolting (n=15). The experiment was repeated thrice. Error bars indicate the standard error (± SE) of three independent experiments. Asterisks indicate significant differences (*$p \leq 0.05$, **$p \leq 0.01$, ***$p \leq 0.001$; unpaired two tailed *t*-test). Scale bar=1cm in (**C**).

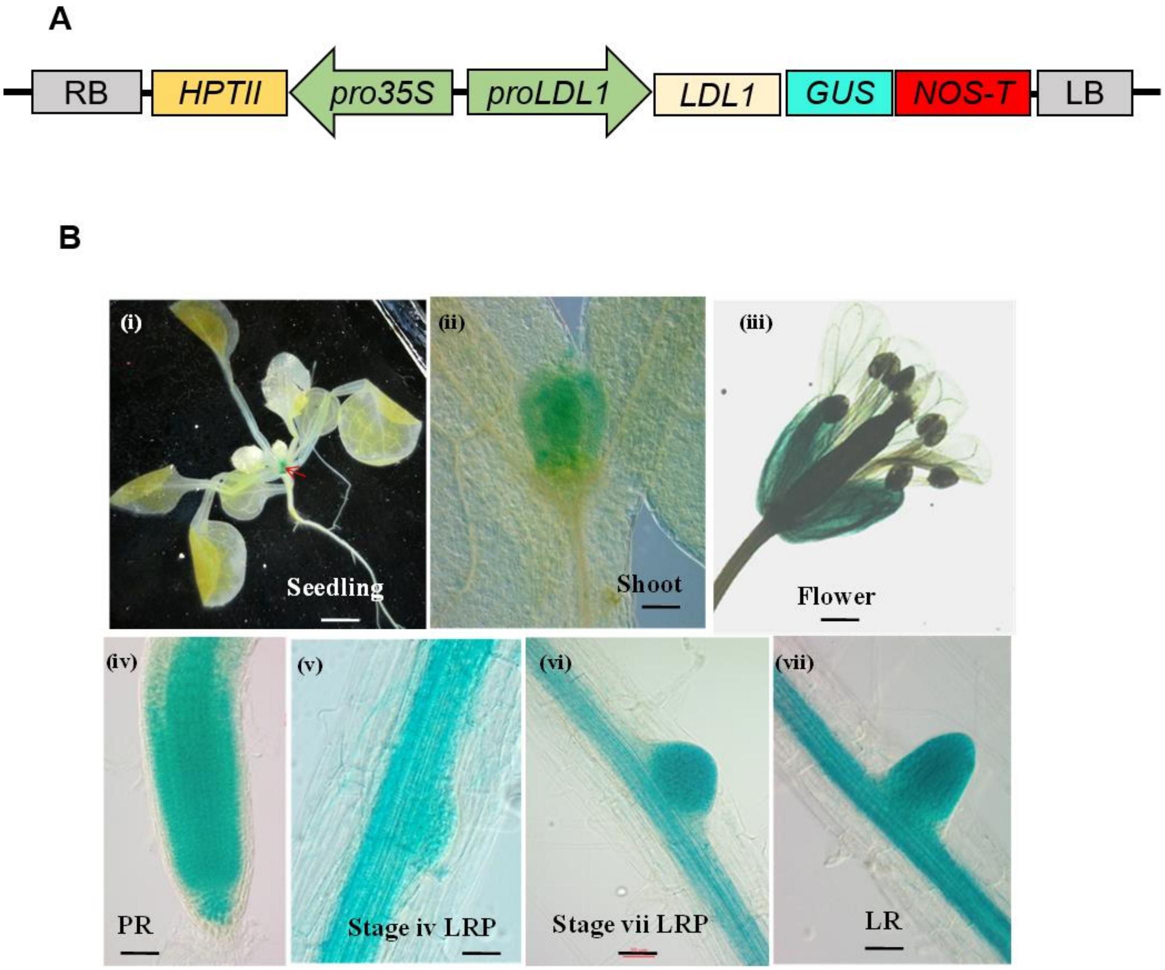


Supplementary Figure 4. **Tissue specific expression pattern of LDL1.** (**A**) Construct showing translational fusion map of *LDL1*. **(B)** GUS activity was observed in different parts of *pLDL1:LDL1-GUS (ldl1)* transgenic line. (i) SAM of four days old seedling (arrowhead indicates SAM) (ii) young leaves, (iii) flowers, (iv) primary root, (v) stage iv LRP, (vi) stage vii LRP, (vii) LR. Scale bar 2 mm in (i) and (iii) and 50 µm in (ii), (iv) – (vii).

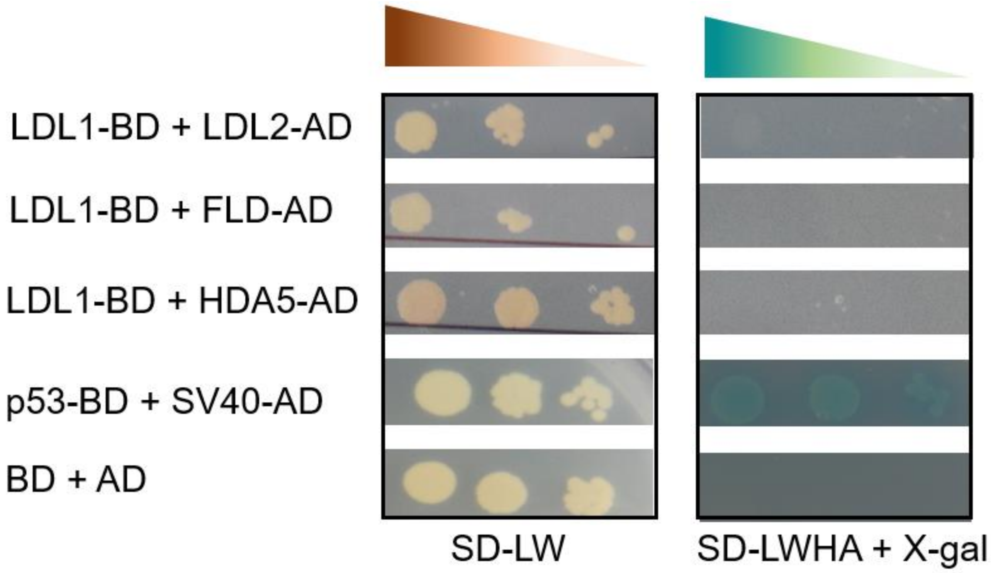


Supplementary Figure 5**. LDL1 doesn't interacts with LDL2, FLD and HDA5 in Y2H assay.** Co-transformed yeast cells were grown on SD-LW and the interaction was detected by the growth of yeast cells on SD-LWHA+X-gal medium and the blue colour indicating MEL1 protein activity. p53-BD and SV40-AD were taken as positive controls and empty vectors were taken as negative controls.

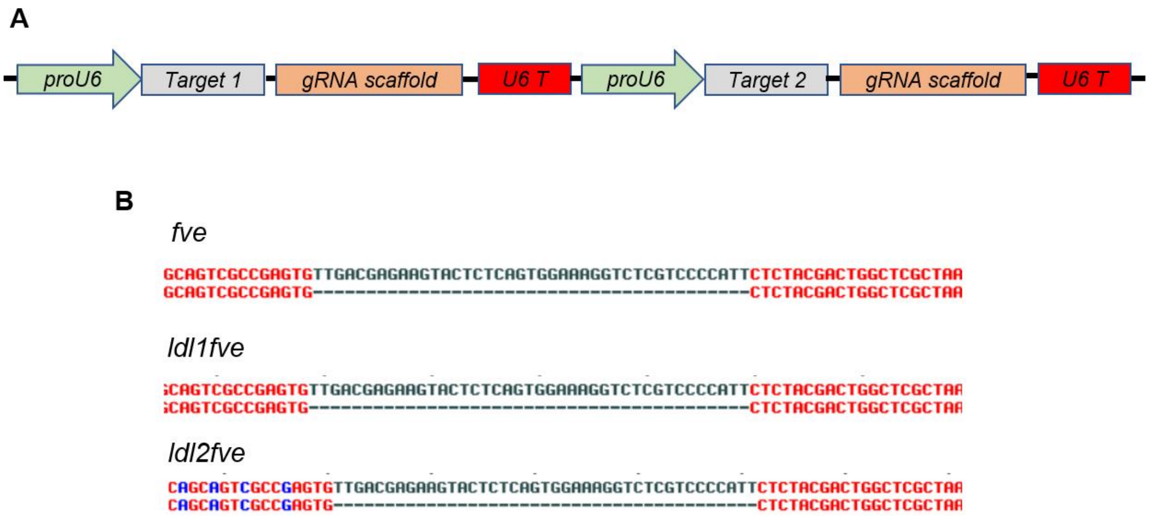


Supplementary Figure 6. **CRISPR/Cas9 mediated mutagenesis in *FVE*** (**A**) Vector map of CRISPR construct. (**B**) Sequencing alignment of gRNA target region showing deletion in *FVE* in the WT, *ldl1* and *ldl2* background**.**

| S.N. | Proteins | WD40 domain repeats | Different complexes |
|---|---|---|---|
| 1 | *At*MSI1 | 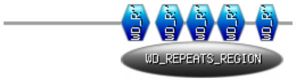 | Chromatin assembly factor 1 (CAF1) [59], Polycomb repressive complex 2 (PRC2) [60]. Interacts with Histone deacetylase 6 [61] |
| 2 | *At*MSI2 | 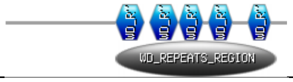 | NA |
| 3 | *At*MSI3 | 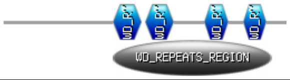 | Imitation Switch (ISWI) chromatin remodeling complex [62] |
| 4 | *At*MSI4/FVE | 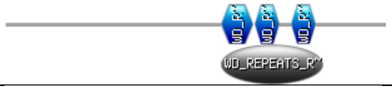 | PRC2 [63], HDA5/HDA6-corepressor complex[25, 64] |
| 5 | *At*MSI5 | 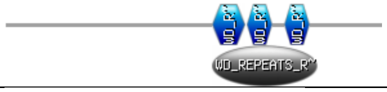 | Interacts with HDA6 [26] |
| 6 | *Sc*MSI1 | 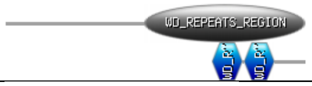 | CAF1[65] |
| 7 | *Sc*HAT2 | 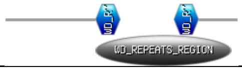 | Histone acetyltransferse (HAT1) [66] |
| 8 | *Dm*Nurf55/p55 | 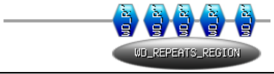 | PRC2[67], HAT1 [68], CAF1 [68] |
| 9 | *Hs*RBAP46 | 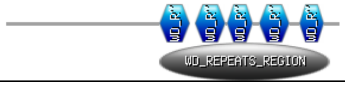 | Histone deacetylation complex (Sin3A)[69], Nucleosome remodeling factor (NURF)[70], Nucleosome remodeling deacetylase (NURD)[71], Polycomb repressive complex 2 (PRC2)[72], Dp, Rb, MuvB (DRM complex), HAT1 [73] |
| 10 | *Hs*RBAP48 | 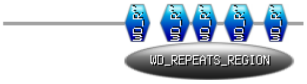 | Sin3A[69], NURF [70], NURD[71], PRC2 [72], DRM complex, CAF1 complex [74] |

Supplementary Table 1. Summary of FVE homologs as a part of different chromatin modifying complexes in *Arabidopsis thaliana* (*At*), *Saccharomyces cerevisiae* (*Sc*), *Drosophila melanogaster* (*Ds*) and *Homo Sapiens* (*Hs*)

Supplementary Table 2. List of primers used in the study

| S.No. | Gene | Sequence | Purpose |
|---|---|---|---|
| 1 | LDL1 F | AAAGGTGGTGATGGTGTTGAG | genotyping |
| 2 | LDL1 R | ATTCCACTTAAGAAGGCTCCG | genotyping |
| 3 | LDL1 F | TCTTTGCACGGTTCCATTAG | qRT PCR |
| 4 | LDL1 R | CTCTTCGCCCCAGAAGTTAC | qRT PCR |
| 5 | LDL1 F | CCCGGGTATGTCAACAGAGACTAAAG | pGEX-4T1 |
| 6 | LDL1 R | CTCGAGCTAATCAAAGATCTGTCG | pGEX-4T1 |
| 7 | LDL1 F | GCGTATTGGGATCAAcATcATCCG | *mLDL1* |
| 8 | LDL1 R | CGGATgATgTTGATCCCAATACGC | *mLDL1* |
| 9 | proLDL1 | AGGTCGACGGCTTCACTAAGGGATTCGTTG | pCAMBIA1301 |
| 10 | LDL1 R | AGCCATGGCTACCGACATTGCGATTAGCG | pCAMBIA1301 |
| 11 | FLC F | AGGCACGACTTTGGTAACACCT | genotyping |
| 12 | FLC R | AAAGGGGGAACAAATGAAAACC | genotyping |
| 13 | FLC F | TGTGGATAGCAAGCTTGTGG | qRT PCR |
| 14 | FLC R | TGAGTTCGGTCTTCTTGGCT | qRT PCR |
| 15 | MAF4 F | AGTACCAGCCGAGCTACCAAG | genotyping |
| 16 | MAF4 R | GTCTTGGGATTTCAAGCCATC | genotyping |
| 17 | MAF5 F | CAATAGATCACGGACATCACG | genotyping |
| 18 | MAF5 R | CCATGCTGGAGAGAGATGAAG | genotyping |
| 19 | MAF4 F | AGACCCATCAAGAGAAGGAGAAGC | qRT PCR |
| 20 | MAF4 R | AGCTGGTTAGTCAAACTCTGGTTC | qRT PCR |
| 21 | MAF5 F | GTCTCTTCCACCGGCAAACTCTAC | qRT PCR |
| 22 | MAF5 R | GATGATCTTGGCCATGCTGTCG | qRT PCR |
| 23 | FT F | CAAGTCCTAGCAACCCTCACC | qRT PCR |
| 24 | FT R | ACGACACGATGAATTCCTGC | qRT PCR |
| 25 | SOC1 F | ATTCGCCAGCTCCAATATG | qRT PCR |
| 26 | SOC1 R | GCCTTCTCCCAAGAGTTTACG | qRT PCR |
| 27 | proMAF4 F | TCTAGATCACCATCATCAACGGCTG | pCAMBIA1304 |
| 28 | proMAF4 R | GGCTGTTTTCTCCGACTAATTCC | pCAMBIA1304 |
| 29 | proMAF4 F | CTGCAGCATAGAAACTCTACTAATAC | pCAMBIA1304 |
| 30 | proMAF4 R | CCATGGCTTTGAACTATTTCTAGT | pCAMBIA1304 |
| 31 | LDL1 F | CACCATGTCAACAGAGACTAAAG | pGBKT7g, pGADT7g |
| 32 | LDL1 R | ATCAAAGATCTGTCGATTCAGTC | pGBKT7g, pGADT7g |
| 33 | LDL2 F | CACCAGAAATGAATTCTCCG | pGBKT7g, pGADT7g |

| | | | |
|---|---|---|---|
| 34 | LDL2 R | ATTAAAATGCAGGGGGTTTAAG | pGBKT7g, pGADT7g |
| 35 | FLD F | CACCATGGTCTCATTCTCCGC | pGBKT7g, pGADT7g |
| 36 | FLD R | TTGTTCAATTTTTTCATCGTCTCACC | pGBKT7g, pGADT7g |
| 37 | HDA5 F | ATCGATGGCTATGGCCGGAGA | pGBKT7g, pGADT7g |
| 38 | HDA5 R | CTCGAGAGGTTTAGACACAGCCTTGTC | pGBKT7g, pGADT7g |
| 39 | FVE F | CACCATGGAGAGCGACGAAGCA | pGBKT7g, pGADT7g |
| 40 | FVE R | AGGCTTGGAGGCACAAGTCATAAC | pGBKT7g, pGADT7g |
| 41 | FVE F | ATATATGGTCTCGATTGCGAGCCAGTCGTAGAGAATGTT | CRISPR |
| 42 | FVE F | TGCGAGCCAGTCGTAGAGAATGTTTTAGAGCTAGAAATAGC | CRISPR |
| 43 | FVE R | AACAGTGTTGACGAGAAGTACTCAATCTCTTAGTCGACTCTAC | CRISPR |
| 44 | FVE R | ATTATTGGTCTCGAAACAGTGTTGACGAGAAGTACTCAA | CRISPR |
| 45 | MAF5 F | CACCATGGATCTTGAAGACAAAACT | pGBKT7g, pGADT7g, |
| 46 | MAF5R | CTTGAGAAGCGGGAGAGTCTC | pGBKT7g, pGADT7g, |
| 47 | FLC F | CACCATGGGAAGAAAAAAACTAGA | pGBKT7g, pGADT7g, |
| 48 | FLC R | ATTAAGTAGTGGGAGAGTCACCGG | pGBKT7g, pGADT7g, |
| 49 | SVP F | CACCATGGCGAGAGAAAAGATTC | pGBKT7g, pGADT7g, |
| 50 | SVP R | ACCACCATACGGTAAGCCGA | pGBKT7g, pGADT7g, |
| 51 | proFT F | GGGCCCAAGTGCATATAGCAGATTTC | pGREENII0800 |
| 52 | proFT R | CTCGAGCTTTGATCTTGAACAAACAG | pGREENII0800 |
